# ssDNA phage FLiP resides in dsDNA form in resistant *Flavobacterium* host

**DOI:** 10.1101/2025.09.12.675792

**Authors:** Kati Mäkelä, Reetta Penttinen, Janne Ravantti, Elina Laanto, Lotta-Riina Sundberg

## Abstract

Most of the environmental flavobacteria decompose organic matter, playing a crucial role in ecosystem balance. Phages infecting these bacteria regulate the host populations and thereby the ecosystem functions, however, bacterial resistance against phage may cause changes in bacterial phenotypic characteristics. ssDNA phage *Finnlakevirus FLiP* infects three known environmental *Flavobacterium* sp. isolates: B330, B167, and B114. Building on our previous FLiP-host interaction studies, we aimed to broaden our understanding of the host perspective by exploring phage resistance mechanisms against FLiP and similar phages. We compared growth dynamics of ancestral and resistant variants to detect the effect of resistance on host fitness. Genomic comparison between the variants was used to identify resistance-related mutations. Additionally, we analysed genomic differences among the three host strains, screening for anti-phage systems. Sequence analyses, PCR, and nuclease treatments of resistant host genomes and plasmids were used to assess the possibility of lysogeny and superinfection immunity. No single mutation or anti-phage system consistently explained FLiP resistance, as these varied across hosts. Resistant B114 contained FLiP genome in circular extrachromosomal dsDNA form, suggesting possible lysogeny. Surprisingly, we found that low quantities of FLiP sequences exist in bacterial populations not exposed to FLiP in the laboratory. This suggests that residing as extrachromosomal dsDNA elements in a small percentage of host cells may be a natural strategy for FLiP type phages to endure unfavourable conditions.

## Introduction

Although the role of viruses is increasingly acknowledged in microbial ecology, due to medical interest, most of the research of phage-bacterium interaction has been done with human bacterial pathogens (Gao et al., 2022; Wieczynski et al., 2023). Furthermore, our perspective of the viral world is highly biased towards dsDNA viruses, as most isolated phages are tailed Myo-, Sipho- and Podoviruses belonging to class *Caudoviricetes*. However, recent genomic data indicates that ssDNA viruses are more prevalent in the environment than previously anticipated (Hopkins et al., 2014; Székely and Breitbart, 2016; Tucker et al., 2011).

*Flavobacterium* species (*Bacteroidetes*) are major players in the freshwater bacterioplankton communities, causing bacterial blooms in natural waters (Eiler and Bertilsson, 2007; Nilsson et al., 2020; Williams et al., 2013). They are among the most important decomposers of organic matter in many types of natural aqueous and soil environments (Bernardet and Bowman, 2006). Interestingly, this bacterial group also hosts the first known ssDNA phage with a lipid membrane (Laanto et al., 2017b; Mäntynen et al., 2020). This untypical virus, *Finnlakevirus* FLiP (*Flavobacterium*-infecting lipid containing phage) infects three flavobacterium hosts isolated from boreal freshwaters (Mäkelä et al., 2024). FLiP is a permanent part of the ecosystem of at least one lake, Jyväsjärvi, and seems to be well adapted to the fluctuating conditions prevailing in the freshwaters of Northern Europe (Mäkelä et al., 2025). Our previous studies show that FLiP-host interactions typically occur on surfaces, possibly during early stages of bacterial attachment and biofilm formation (Mäkelä et al., 2024).

In the case of flavobacteria, resistance against dsDNA phages typically cause mutations in the type IX secretions system (T9SS) genes (Kunttu et al., 2021; Laanto et al., 2017a), which are associated with flavobacterial gliding motility machinery (McBride and Nakane, 2015). This results in loss of motility, deficiencies in protease secretion and, in pathogenic flavobacteria, loss of virulence (Barbier et al., 2020; Kharade and McBride, 2015; Laanto et al., 2014; Rhodes et al., 2011). Mutations may also lead to inability of the bacterium to adhere to surfaces (Lasica et al., 2017). However, possible genetic mutations related to resistance against ssDNA phages are so far not known. CRISPR-Cas based immunity as well as superinfection immunity, caused by lysogenic presence of a phage in an infected cell, are among resistance mechanisms that have been found in flavobacteria (de Freitas Almeida et al., 2022; Laanto et al., 2020; Nilsson et al., 2020). FLiP-type prophages have been found from bacterial metagenomes (Yutin et al., 2018), but lysogenic cycle has not been shown for FLiP so far. However, our previous studies suggest that FLiP endures certain conditions such as anoxic and warm periods in association with the host rather than in virion form (Mäkelä et al., 2025).

In this study, we explored the possible mechanisms which *Flavobacterium* sp. strains B330, B167 and B114 may use as protection against infections by FLiP and similar phages. We exposed the three host bacteria to FLiP to select for phage resistant variants. Then, we used genomic sequencing and phenotypic experiments such as growth assays to compare differences between the host strains and between the phage resistant variants and ancestral strains. B330 and B167 were genetically closely related, whereas B114 was observed to be a more distant relative. The phage-exposed resistant variant of B114 was shown to carry FLiP as extrachromosomal dsDNA form, suggesting that lysogeny and superinfection exclusion might provide the phage resistance in this host. We also inferred that phages like FLiP might exist as extrachromosomal dsDNA form in many natural *Flavobacterium* populations, albeit sometimes only in a small fraction of the cells. These dsDNA elements could represent long-lived replicative forms (RF) of delayed infection, pseudolysogenic, or lysogenic states. FLiP-type phages might also flexibly switch between these life cycle options, at least in some hosts, adapting to the most advantageous strategy based on environmental conditions without permanently committing to any single state.

## Methods

### Bacterial strains and resistant variants

*Flavobacterium* sp. strains B114, B167 (Laanto et al., 2011) and B330 (Laanto et al., 2017b), the isolation host of FLiP, were used in this study. Bacterial host variants resistant to FLiP were isolated by first spreading 100 µl high-titer (~10^10^ plaque forming units (PFU) ml^-1^) FLiP lysate on a Shieh (Song et al., 1988) agar [1 % (wt/vol)] plate. After the lysate had absorbed, 100 µl of dilution (10^2^ colony forming units / ml) of overnight (21°C, 120 rpm) grown, wild type host bacterial culture (B330, B167 and B114) was spread on top of the phage lysate. After incubating 5 days at room temperature (RT), the colony morphology of the appeared resistant colonies was recorded. Non-spreading colonies were picked, and three rounds of colony purifications were implemented. To ensure that phage particles were not left as contaminants in the purified variants, liquid cultures were grown overnight (RT, 120 rpm), centrifuged, filtered (0.2 µm), and supernatant spotted onto susceptible hosts in a drop assay.

The purified resistant bacterial variants were tested for resistance to FLiP infections by double layer plaque assay: 3 ml of Shieh soft agar [0,7 % (wt/vol)] was tempered into + 47°C. 100 µl of overnight grown host bacterium and 100 µl of phage dilution were mixed into soft agar and the mixture was poured onto a Shieh agar plate. Plates were incubated 2 days at RT. If no plaques appeared, the host was considered resistant and designated B330r, B167r and B114r.

To record the typical colony morphologies of each ancestral and resistant strain, bacterial dilutions were spread on Shieh-agar plates in triplicates and incubated three days at RT. Plates were imaged with ChemiDoc MP Imaging system (Bio-Rad) using application for Ethidium Bromide Gel (602/50, UV Trans) and Auto Optimal exposure.

### Growth of the ancestral and resistant bacteria

A Bioscreen C (Growth Curves Ltd, Helsinki, Finland) spectrophotometer was used to monitor the growth of the ancestral and phage resistant bacterial variants. Before the experiment the optical densities of the fresh overnight-grown bacterial cultures were adjusted to ~0.20 (dilution of approximately 1:10 o/n culture to fresh Shieh) in A570 to minimize the differences in the initial turbidity between strains. Each strain was transferred (220 µl per well, 5 replicates per treatment) into a BioScreen Honeycomb plate (100-well plate, Oy Growth Curves Ab Ltd). Next, FLiP was added into the host cultures in 20 µl volume (4,4 E+5 PFUs) and corresponding volume of Shieh was added to the no-phage controls. The bacterial growth was measured for 72 hours at 22°C at 1-hour intervals (absorbance at 600 nm).

### Genome sequencing and analyses

DNA was extracted from phage resistant strains B330r, B167r and B114r using Genomic tip 100/G (Qiagen) according to instructions of the manufacturer and using the recommended buffers. DNA pellet from Genomic tip extraction after isopropanol precipitation was resuspended in 120 µl of elution buffer from DNeasy Blood & Tissue kit (Qiagen). DNA from ancestral strains B167 and B114 was extracted using DNeasy Blood & Tissue kit, according to manufacturer instructions and using the elution volume of 100 µl. Ancestral B330 DNA was purified using Genomic-tip 20/G kit (Qiagen), according to manufacturer instructions.

Genomic DNA of the ancestral phage-susceptible strains B114 and B167 was subjected to microbial whole genome library preparation (350bp) and Illumina NovaSeq (150 bp paired end) sequencing in Novogene. B330 was sequenced commercially in Institute of Biotechnology in University of Helsinki using Nextera library preparation to prepare 500-600 bp sequencing library, and producing 300 bp paired end reads with Illumina MiSeq. The genomic DNA of the resistant variants B114r and B167r was sequenced in Novogene with PacBio RSII sequencing. The genomes of B114r and B167r were assembled and assured to produce a circular chromosome by SMRT portal (Version 3.2.0) (Li et al., 2010, 2008). For B330r strain, single-molecule real-time sequencing was performed on a PacBio RSII sequencer (BGI). A draft genome of B330r was assembled with SPAdes 3.15.5 (Antipov et al., 2016) as a hybrid assembly, using both B330 Illumina short reads and B330r PacBio subreads. The overlapping ends were manually trimmed to produce a circular chromosome using Geneious Prime ® 2023.2.1. (Biomatters Ltd.). Subreads of B330r were mapped on the chromosome with minimap2 (v2.24) (Li, 2018) and the final consensus sequence was determined with samtools (v1.16) consensus tool (Li et al., 2009). Finally, the *dnaA* gene was set as the starting point of the B330r genome.

The complete genomes of B114r, B167r and B330r were annotated with the NCBI Prokaryotic Genome Annotation pipeline (Haft et al., 2018; Li et al., 2021; Tatusova et al., 2016) and deposited in the Genbank under BioProject PRJNA947224. Bacterial anti-phage systems were identified with PADLOC v2.0 (Payne et al., 2022, 2021) and prophages detected with VirSorter2 (Guo et al., 2021) and PHASTER (Arndt et al., 2016). The taxonomic classification of the isolated strains B114r, B167r and B330r was performed by first retrieving all the complete *Flavobacterium* genomes (N=69) from GenBank. *Flavobacterium* sp. B183 (GenBank Acc. No. CP097434) and *F. johnsoniae* UW101 (GenBank Acc. No. CP000685) were also included in the analysis as reference. The core genome shared by the collected *Flavobacterium* genomes was assessed with Roary (Page et al., 2015). A phylogenetic tree was built based on the presence of accessory (non-core) genes. For visualization, a final subtree was extracted to include the 29 closest related *Flavobacterium* strains or species.

The relatedness of the host strains B114r, B167r and B330r as well as *Flavobacterium* sp. B183 and *F. johnsoniae* UW101 was assessed by identifying the shared gene content by pairwise all-against-all comparisons of all amino acid sequences. This analysis was conducted using the Needle program from the EMBOSS package (Rice et al., 2000), with a sequence similarity cutoff of 90%.

### Analysis of FLiP-resistant variants

The potential genetic changes conferring resistance against FLiP phage were identified by trimming the ancestral host reads with fastp 0.23.4 (Chen, 2023) and mapping them against the assembled genome of the observed resistant variant using breseq 0.38.1 (Barrick et al., 2014). The tool was run as clonal mode to detect mutations present in all reads.

### Plasmid DNA extractions and analysis of ancestral and resistant bacterial strains

Qiaprep Spin Miniprep Kit (Qiagen) was used to extract plasmid DNA from overnight cultures of ancestral (B330, B167 and B114) and resistant (B330r, B167r and three resistant variants of B114: B114r, B114rB and B114rC) bacterial strains as well as from overnight culture of control bacterium *Flavobacterium* sp. B183 (Laanto et al., 2011), previously observed to carry a plasmid (Sundberg et al., 2016). Genomic DNA of B183 as well as 32 other *Flavobacterium* strains (Supporting information Table S1) was extracted using DNeasy Blood & Tissue kit (Qiagen) according to manufacturer’s instructions.

Genomic and plasmid DNA from B330, B167, B114, B330r, B167r, B114r, B114rB, B114rC and B183 as well as genomic DNA from the above mentioned 32 *Flavobacterium* strains were used as templates in PCR reactions (Supporting information). Primers were used to amplify ~500 bp area of either major capsid protein (MCP) or replication initiator protein (Rep) of FLiP. PCR products were run in 1 % agarose gel.

Plasmid DNA extracted from B114r was digested using restriction enzymes *Bgl*II (Thermo scientific), *Xba*I (Thermo scientific), *Bgl*I (Fermentas) and *Hind*III (Thermo scientific). Digested plasmids and an undigested control plasmid were run in 1 % agarose gel.

To track the potential lysogenic or pseudolysogenic state of FLiP within the resistant host cell, the produced PacBio subreads were mapped against reference FLiP genome (NC_047837) with minimap2 (v2.24). Only primary hits were included, and the matching reads were filtered to include only unique reads which were mapped against the FLiP genome and the coverage of the phage genome was determined with samtools (v1.16). To exclude the potential intracellular carriage of FLiP in the ancestral strains (prior to FLiP exposure under laboratory conditions), we also mapped the trimmed Illumina reads of B114, B167 and B330 on FLiP genome using bowtie v2.5.3 (Langmead and Salzberg, 2012).

## Results

### Colony morphologies of the ancestral and resistant host variants

Ancestral B330 grew on Shieh-agar plates as thin, effectively spreading, pale-yellow colonies (Figure 1). Colony morphology of ancestral B167 was highly similar to that of B330, but slightly less spreading and more intensive in colour. Colonies of ancestral B114 were intensely yellow and markedly thicker and less spreading than those of B330 or B167. There were a few non-spreading, sharp-edged B114 colonies (~12 %), which were smaller in diameter compared to the spreading ones.

**Figure 1.**
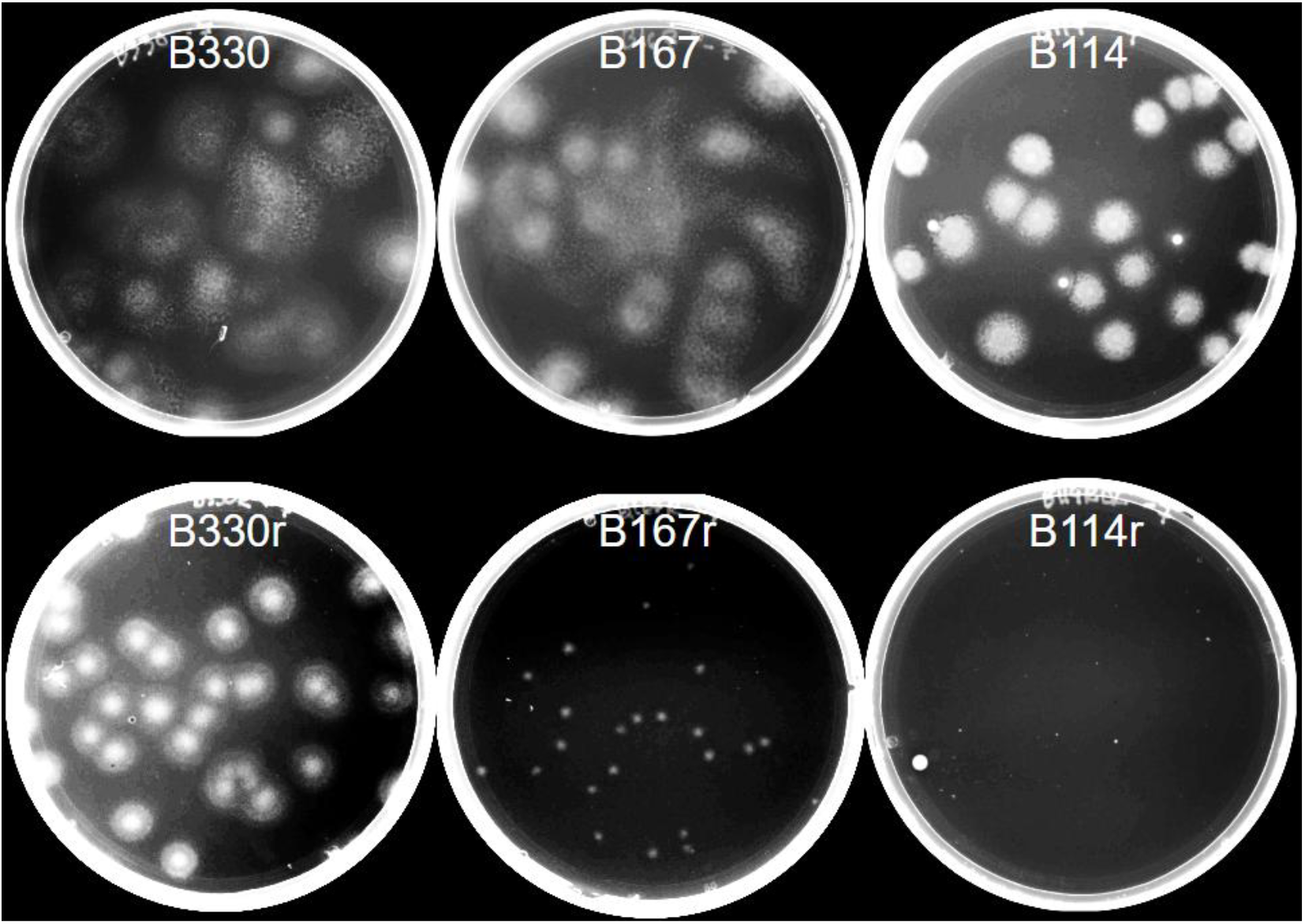
Colonies of the ancestral host strains of FLiP, Flavobacterium sp. B330, B167 and B114, and the three times purified phage resistant variants B330r, B167r and B114r, on Shieh-agar plates after three days of incubation at room temperature. Plates were imaged with ChemiDoc MP Imaging system (Bio-Rad). Figure is composed of separate plate images. Brightness and contrast have been adjusted for better visibility of colonies.

A change in the colony morphology was observed in all the host bacteria exposed to FLiP. The phage resistant B330 and B167 colonies, which formed when the ancestral strain was exposed to FLiP, were either non-spreading or only moderately spreading while all the B114 resistant colonies were non-spreading with sharp edges. One three times purified, resistant variant with non-spreading colony morphology of strains B330 and B167 was chosen for further analyses and named B330r and B167r. Three thrice purified resistant colonies of B114 were chosen for further analyses and named B114r, B114rB and B114rC. Results of the sequenced variant, B114r, are presented below since all B114 resistant variants were seen to produce identical results.

When dilutions of B330r, B167r and B114r were plated on Shieh-agar plates, all resistant variants formed smaller colonies compared to the respective ancestral variants (Figure 1). B114r colonies were either approximately the same size as the non-spreading colonies of ancestral strain or extremely small. All B114r colonies had sharp edges, while B167r colonies were small, but had smoother edges (Figure 1). B330r colonies were smaller and thicker than the ancestral ones but still retained smooth edges. B330r colonies were larger compared to colonies of other resistant variants.

### Effect of phage resistance on host growth in liquid

Respective ancestral and resistant variants of each bacterial strain grew similarly in liquid, when phage was not present (Figure 2). In the presence of FLiP, the growth (i.e. the population density at 48 h) of the resistant variants was significantly higher than the susceptible ancestral strains (Mann-Whitney tests, p < 0.05 for all comparisons). The difference was largest in B114r compared to B114 (Figure 2). In general, growth of *Flavobacterium* sp. B330 and B167 was higher than that of B114.

**Figure 2.**
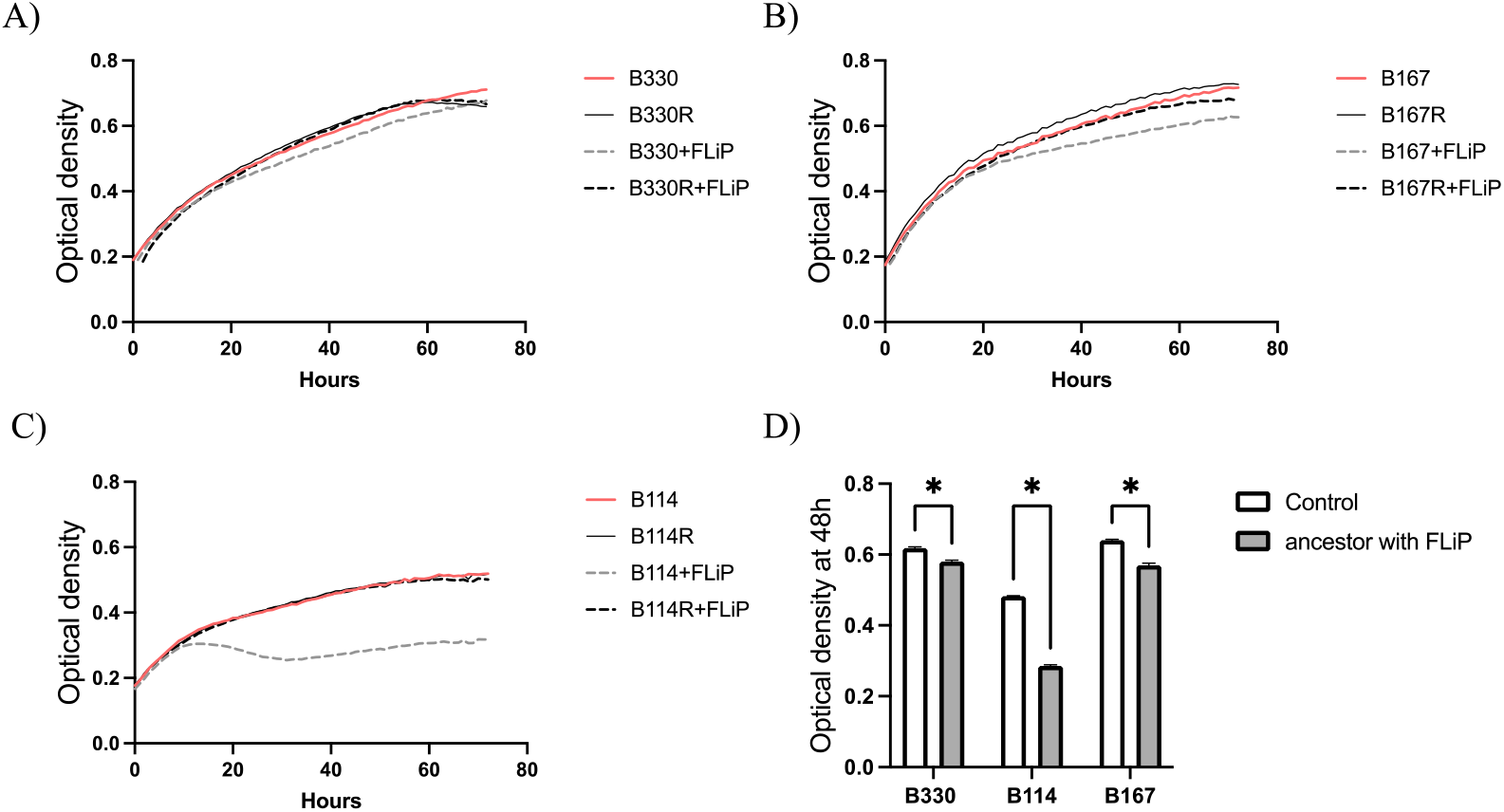
Mean growth (n=5) of ancestral Flavobacterium sp. A) B330, B) B167 and C) B114 strains compared to the phage resistant variants B330r, B114r and B167r with and without presence of FLiP. D) Bacterial population density of ancestral strains (B330, B114 and B167) at 48h with and without FLiP. Asterisk indicates statistical significance (p< 0.05). Optical density (OD at 600 nm) was measured in 1-hour intervals for 72 hours.

### Genomic comparison of *Flavobacterium* hosts

Genomes of the ancestral host strains of FLiP, *Flavobacterium* sp. B330, B167 and B114, were compared to their FLiP-resistant variants B330r, B167r and B114r. Genomic mutations were found in all strains (Table 1, Supplementary Table 1). However, these mutations concerned different protein-coding sites in each strain. Large deletions or mutations were not observed. Interestingly, the less-spreading colony morphology of the resistant variants (Figure 1) seemed not to associate with mutations in T9SS-related genes. In B114, multiple short (1-2 bp) insertions or deletions were found in gene cluster coding for RagB/SusD family nutrient uptake outer membrane protein (Table 1). In case of B167 mutations were in a gene involved in cysteine biosynthesis and in a large cluster of regulatory, signal transduction and sensory proteins. Observed mutations in B330 included AarF/UbiB family protein and genes contributing to formation of amyloid fibers called curli and in a gene oxidizing D-2-hydroxyacids to 2-oxoacids.

**Table 1.**
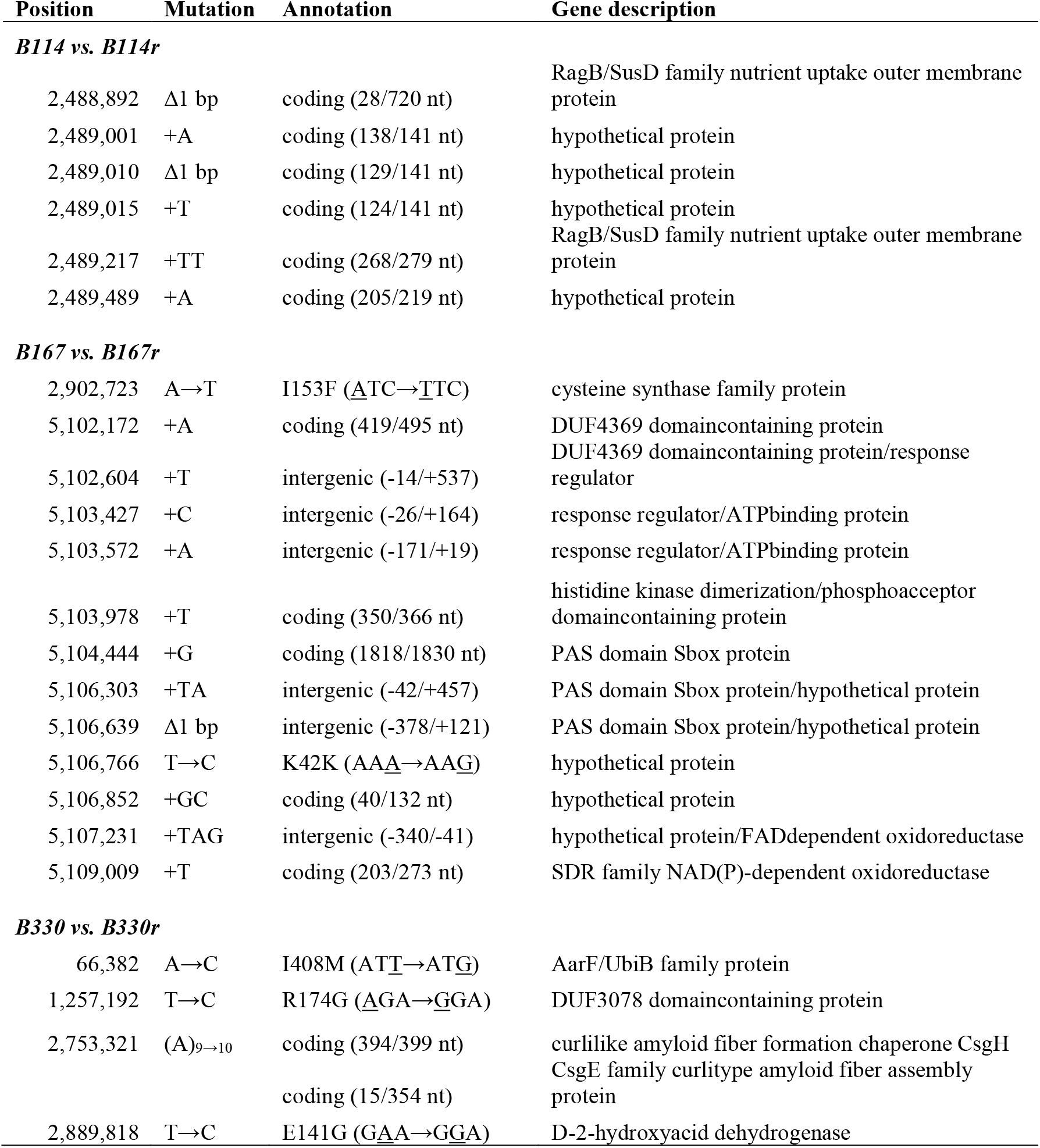
Identified mutations in FLiP-resistant variants.

In the further analyses, the complete genomes of resistant bacterial variants sequenced with PacBio were used. The genomes of the hosts of FLiP were 5.4-5.8 Mbp long, with GC% of 34.2 (B114r), 33.7 (B167r) and 33.9 (B330r) (Table 2A). The genomes of B114r, B167r and B330r were screened for the presence of anti-phage systems. These bacterial strains do not have CRISPR-Cas systems, but a selection of other anti-phage systems was found that were mostly distinct to each bacterial strain (Table 2B, Supplementary table 2). SoFic anti-phage system is the only system shared by all the three host strains of FLiP (Table 2B). Restriction-modification (RM) type I and type IV systems, which detect and cleave unmethylated or modified dsDNA respectively (Dryden et al., 2001; Loenen and Raleigh, 2014), as well as Septu type I system, which causes DNA or RNA damage by using ATPase and HNH nuclease (Doron et al., 2018; Wu et al., 2024), were shared by both B114r and B330r. Both B167r and B330r have a VSPR system. In B114r several antiphage systems cluster together (4 604 351 – 4 646 618), forming a defense island (Supporting information Figure S1) including mobile genetic elements (Table 2, Supplementary Table 2).

**Table 2.**
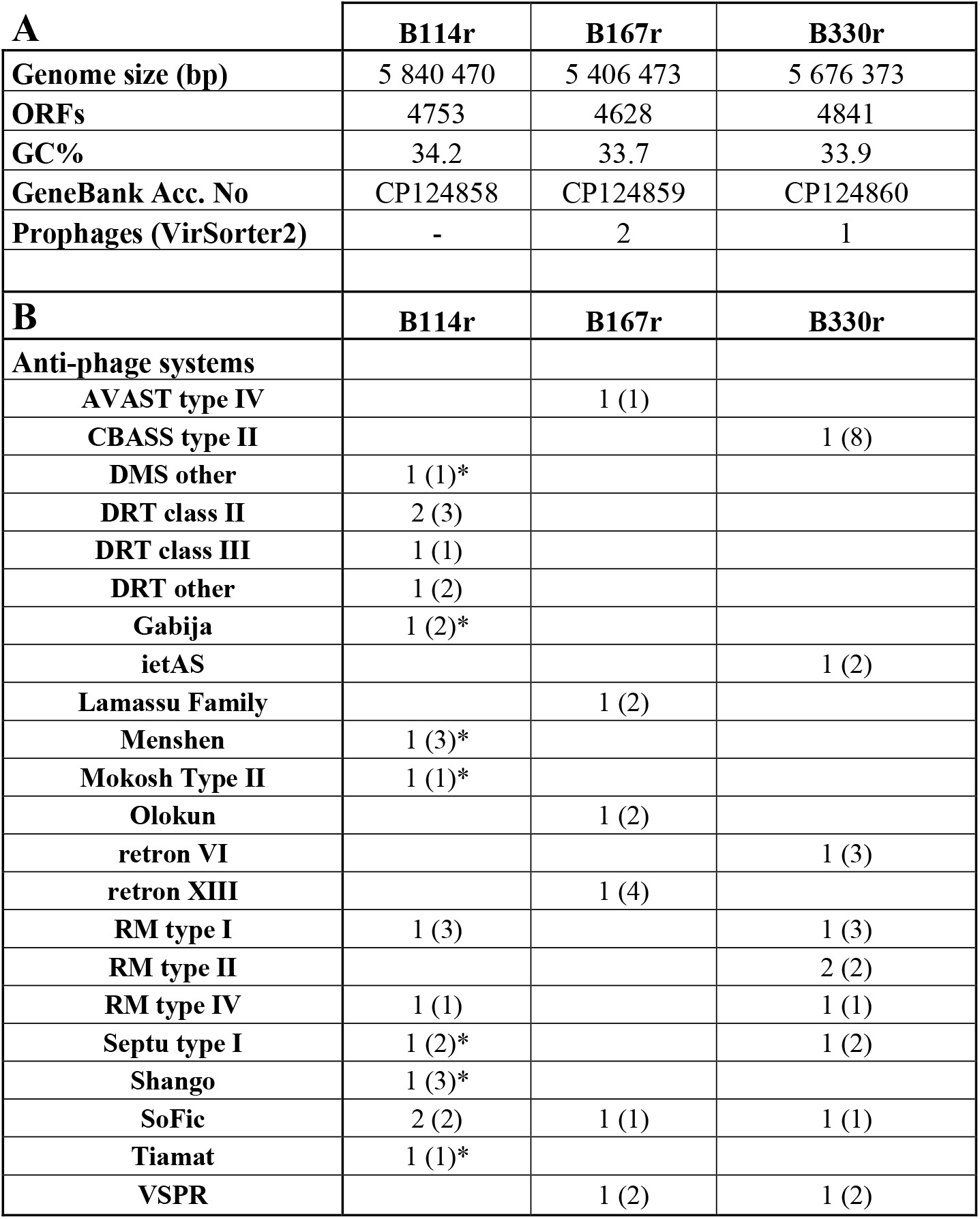
A) Features of Flavobacterium sp. genomes and B) complete anti-phage systems and associated genes (in parentheses) detected with Padloc v2.0. Asterisk indicates antiphage systems that cluster together as a defense island in B114r.

| **A** | **B114r** | **B167r** | **B330r** |
| --- | --- | --- | --- |
| **Genome size (bp)** | 5 840 470 | 5 406 473 | 5 676 373 |
| **ORFs** | 4753 | 4628 | 4841 |
| **GC%** | 34.2 | 33.7 | 33.9 |
| **GeneBank Acc. No** | CP124858 | CP124859 | CP124860 |
| **Prophages (VirSorter2)** | - | 2 | 1 |
| **B** | **B114r** | **B167r** | **B330r** |
| **Anti-phage systems** |  |  |  |
| AVAST type IV |  | 1 (1) |  |
| CBASS type II |  |  | 1 (8) |
| DMS other | 1 (1)* |  |  |
| DRT class II | 2 (3) |  |  |
| DRT class III | 1 (1) |  |  |
| DRT other | 1 (2) |  |  |
| Gabija | 1 (2)* |  |  |
| ietAS |  |  | 1 (2) |
| Lamassu Family |  | 1 (2) |  |
| Menshen | 1 (3)* |  |  |
| Mokosh Type II | 1 (1)* |  |  |
| Olokun |  | 1 (2) |  |
| retron VI |  |  | 1 (3) |
| retron XIII |  | 1 (4) |  |
| RM type I | 1 (3) |  | 1 (3) |
| RM type II |  |  | 2 (2) |
| RM type IV | 1 (1) |  | 1 (1) |
| Septu type I | 1 (2)* |  | 1 (2) |
| Shango | 1 (3)* |  |  |
| SoFic | 2 (2) | 1 (1) | 1 (1) |
| Tiamat | 1 (1)* |  |  |
| VSPR |  | 1 (2) | 1 (2) |

Relatedness between the host strains (Table 3) and within related *Flavobacterium* species was studied by determining the pangenome and constructing a phylogenetic tree based on non-core genes (Figure 3). B330r and B167r were closely related to each other, while B114r was more distant (Figure 3). *Flavobacterium* sp. B183, which was used in some experiments as a control, was seen to be closely related to B114r. One of the most extensively studied strains within the *Flavobacterium* genus. *F. johnsoniae* UW101, was found to be the most closely related to B330.

**Table 3.** Number of shared genes between the Flavobacterium whole genomes (excluding pseudogenes). The number of homologous genes for each pairwise genome comparison is presented with color-coding with a gradient from blue to red; red indicating the pair with the highest number of shared genes and blue indicating the pair with the fewest shared genes. Genes with ≥ 90% amino acid sequence similarity were considered shared. The total number of genes in each genome is shown in parentheses.

| **Genome (genes)** | **UW101(5017)** | **B183(4406)** | **B114r (4712)** | **B330r (4821)** |
| --- | --- | --- | --- | --- |
| B183 (4406) | 991 |  |  |  |
| B114r (4712) | 1131 | 1898 |  |  |
| B330r (4821) | 1711 | 635 | 693 |  |
| B167r (4613) | 1322 | 553 | 647 | 3041 |

**Figure 3.**
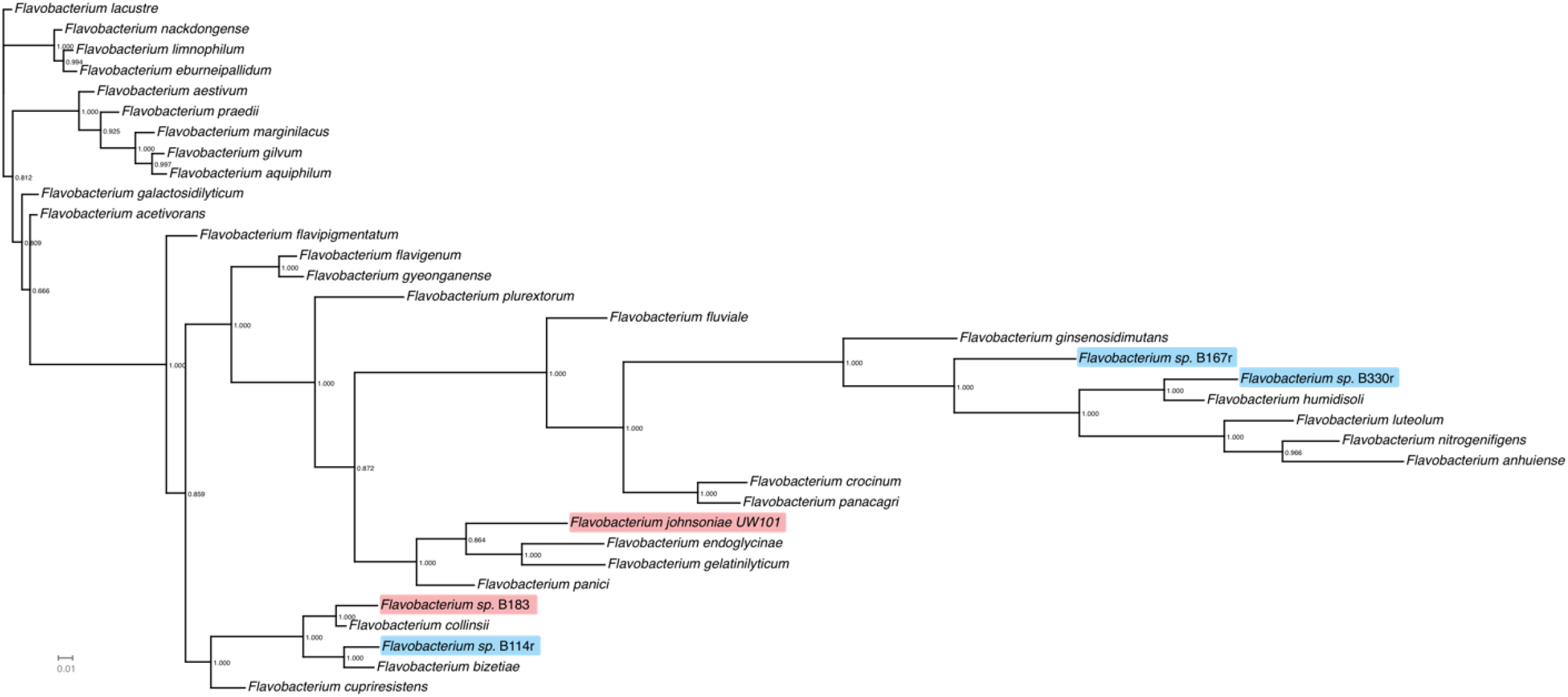
Subtree of Flavobacterium species based on the presence of accessory genes. FLiP host strains are shown in blue, and the control strains used in this study in pink.

Number of shared genes (minimum 90% amino acid sequence similarity) was detected with pairwise comparisons of the *Flavobacterium* sp. strains B114r, B167r, B330r and B183 and *F. johnsoniae* UW101 (Table 3). Shared genes between the strains confirmed the relatedness patterns of FLiP hosts seen in the phylogenetic tree (Figure 3), with B330r and B167r sharing the highest number of similar genes (> 3000), whereas the number of shared genes between B330r and B114r was only 693. B114r and B183 had almost 1900 similar genes. Genetic similarity of hosts of FLiP with *F. johnsoniae* UW101 was highest in B330r with 1711 shared genes.

### Analysis of the presence of FLiP in the phage-resistant variants

To analyze the presence of FLiP DNA in the ancestral and phage-exposed bacterial hosts, we extracted both genomic and plasmid DNA from all strains and used PCR targeting either FLiP replication initiation protein (Rep) or major capsid protein (MCP) encoding genes. Plasmid DNA extracted from the ancestral or resistant FLiP host strains or from the control bacterium B183 produced a PCR product (Figure 4). The strongest bands in agarose gel were observed in FLiP-exposed B114 variants B114r, B114rB and B114rC. Also, the resistant variants B167r and B330r as well as B183 produced stronger bands compared to ancestral host strains B330, B167 and B114. When bacterial genomic DNA was subjected to PCR with above mentioned primers, the result was almost similar, but ancestral B167 or B114 strains did not produce visible signals in agarose gel (Figure 4).

**Figure 4.**
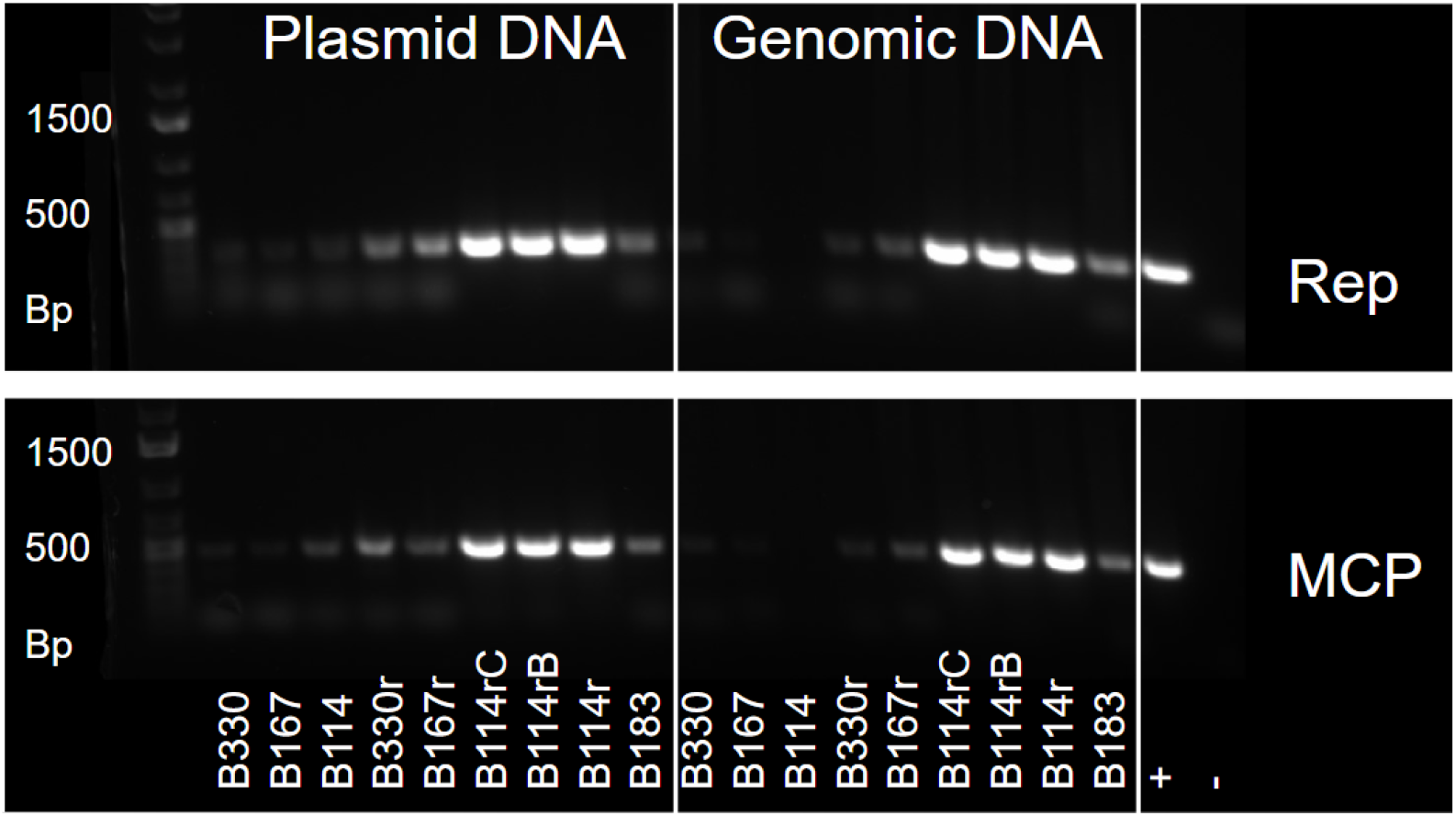
Plasmid and genomic DNA extracted from ancestral and resistant host strains of FLiP subjected to PCR with FLiP-specific primers targeting replication initiation (Rep) and major capsid protein (MCP) genes.

Based on this result, we explored the possibility of FLiP presence in other *Flavobacterium* hosts. The genomic DNA of 32 *Flavobacterium* strains, other than FLiP hosts, were used in PCR reactions (Supporting information Table S1) with FLiP MCP or Rep primers. Seven strains (B067, B138, B169, B207, B222, B350 and B480) out of the 32 tested yielded positive PCR results when using MCP primers, whereas all other PCR reactions, including all of those with Rep primers, were negative (Table S1). Notably, some of the PCR products generated by the FLiP MCP primers for these 7 strains differed in size from those produced by FLiP hosts and B183. These results indicate that B067, B138, B169, B207, B222, B350 or B480 do not harbor FLiP as a prophage but may contain other prophages with MCP sequences similar to those of FLiP.

Further analyses on plasmid DNA extracted from B330r, B167R and B114r were made to confirm that the positive results in PCR indeed originate from presence of FLiP phage in the resistant bacterial hosts and not from other related prophages or the host genome. It was also important to preclude the possibility of particle contamination of resistant strains, which in case of FLiP was possible by distinguishing between ssDNA and dsDNA.

First, in order to identify the existence of FLiP genome within the resistant host cells, we explored the subreads from PacBio sequencing to detect reads matching FLiP genome. We were able to identify the FLiP genome from B114r with a 206x sequencing depth, whereas no matching reads were found from B167r or B330r. None of the ancestral (B114, B167, B330) short reads matched FLiP genome sequence, confirming that these strains did not readily carry FLiP-like phage as lysogenic nor pseudolysogenic form prior to the phage exposure.

Next, the plasmid DNA extracted from the sequenced resistant variant, B114r, was digested with four different restriction enzymes which cleave only dsDNA in sequence specific sites.

*Bgl*II and *Xba*l both have one target site in FLiP genome whereas *Bgl*I is supposed to cleave it in two different sites, and *Hind*III in five different sites. Undigested FLiP plasmid was seen as a ~5500 bp band on agarose gel (Figure 5), which refers to the tendency of circular plasmids to take a supercoiled conformation. As plasmid DNA was cleaved in one site by *Bgl*II and *Xba*l, the secondary structures were relaxed, and the band seen in agarose gel was ~9000 bp equating well the genome size of FLiP (9174 nt) (Figure 5). Based on FLiP genomic sequence, *Bgl*I cleaves FLiP genome as dsDNA plasmid in two sites producing two fragments sized 1865 ja 7309 bp. *Hind*III should cleave the plasmid into 3541, 2717, 1699 ja 1169 and 48 bp fragments. Both *Bgl*I and *Hind*III digestions produced exactly the expected results except for the very small 48 bp band which could not be seen in gel (Figure 5). Taken together, restriction analysis confirmed that FLiP exists in B114r as circular dsDNA elements.

**Figure 5.**
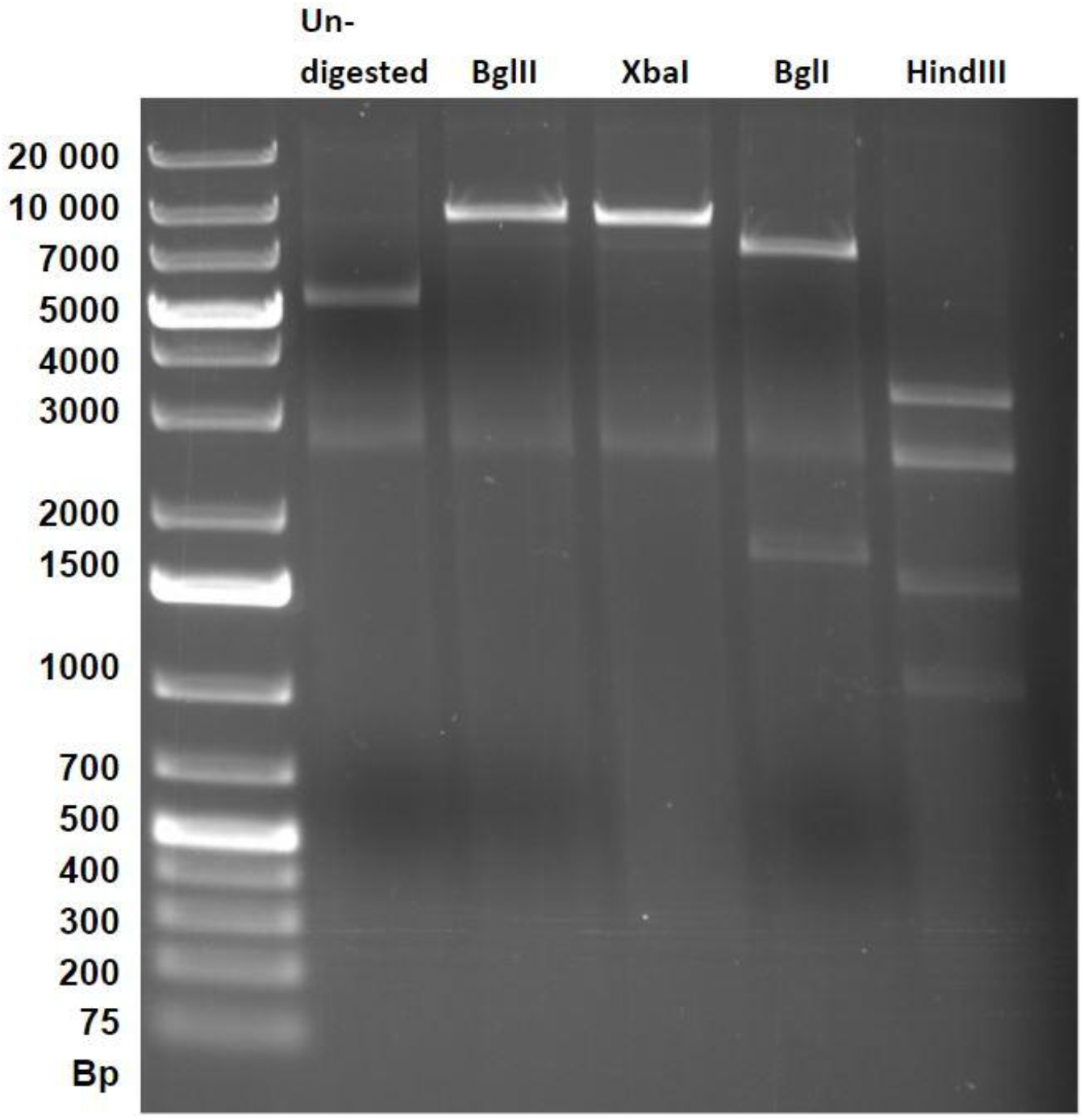
B114r plasmid DNA digested with restriction enzymes targeting sequence specific sites of dsDNA.

## Discussion

Phage resistance often causes fitness costs for the bacterium, which leads to coexistence of both resistant and susceptible variants in bacterial populations (Hasan and Ahn, 2022; Oechslin, 2018). However, the influence of phage resistance against ssDNA phages on bacterial host fitness is not well understood. Our results indicate that at least a subset of *Flavobacterium* population can either become resistant to FLiP relatively easily or some of the cells in a population are constantly resistant. We could not detect differences between the growth of ancestral and FLiP-resistant host variants in liquid cultures without the presence of FLiP, but possible costs may be condition-dependent, since mutations influencing bacterial nutrient intake and metabolism were identified (see below).

Bacterial defence systems, comprising of one or several genes, as well as accordant phage anti-defence mechanisms, are known for many types of phage-bacterium pairs both in clinical settings and in nature (Bleriot et al., 2024; Egido et al., 2022; Hasan and Ahn, 2022; Labrie et al., 2010; Skanata and Kussell, 2021). Here, we investigated three different *Flavobacterium* hosts of FLiP. Each of them had a different set of anti-phage systems, yet it remains to be evaluated which of them, if any, are used against FLiP. SoFic anti-phage system was the only system shared by all the three host strains. This system suppresses phage replication most likely by inactivating phage proteins by AMPylation, and it is one of the most prevalent defense systems worldwide in bacterial metagenomes (Beavogui et al., 2024; Millman et al., 2022). The wide variety of anti-phage systems found in the genomes of the three *Flavobacterium* strains indicates that a diverse arsenal of resistance mechanisms may be needed to survive under variable prevailing environmental conditions. It is also possible for bacteria to use several anti-phage systems concurrently, and in some cases domains of incomplete defence systems may be utilized by the active ones (Wu et al., 2024).

Genetic differences between the ancestral and the resistant variants of hosts of FLiP were found, but they did not occur in genes encoding for known anti-phage systems. Nor could we identify common mutations in the FLiP-resistant bacterial hosts. In B167 and B114, the phage resistance caused a clear change in bacterial colony morphology. In previous studies on *Flavobacterium*-phage interactions using tailed phages, resistance typically selects for mutations in T9SS and gliding motility genes (Castillo et al., 2021; Kunttu et al., 2021; Shrivastava et al., 2013), which were not observed here in the FLiP-resistant hosts. In B114, multiple short (1-2 bp) insertions or deletions were found in the gene cluster coding for nutrient uptake outer membrane protein (Table 1). RagB/SusD family nutrient uptake proteins contribute to binding and uptake of carbohydrates in *Bacteroidetes* species, and they are typically part of a gene cluster known as bacterial polysaccharide utilization locus (Koropatkin et al., 2008; Terrapon et al., 2015). In case of B167, mutations were in a gene involved in cysteine biosynthesis and in a large cluster of regulatory, signal transduction and sensory proteins such as response regulators, histidine kinase (Wolanin et al., 2002) and PAS domain S-box proteins (Taylor and Zhulin, 1999), suggesting this cluster may be involved in environmental sensing and cellular response mechanisms. Indeed, we have previously shown that conditions such as temperature and structure of the environment strongly affect both the growth of the host bacterium and the FLiP-host interactions (Mäkelä et al., 2024). The observed mutations in B330 may have effects on coenzyme Q biosynthesis and its cellular distribution as an AarF/UbiB family protein was mutated (Kemmerer et al., 2021; Poon et al., 2000). There were also mutations in B330 genes contributing to formation of amyloid fibers and consequently bacterial attachment to surfaces and biofilm formation (Barnhart and Chapman, 2006) as well as in a gene oxidizing D-2-hydroxyacids to 2-oxoacids, potentially involved in various metabolic pathways (Matelska et al., 2018). In conclusion, mutations in genes and gene clusters encoding for proteins involved in environmental sensing, signal-transduction, cell adhesion to surfaces, nutrient uptake and utilization processes, biosynthesis and regulatory functions may contribute to the observed changes in colony morphologies, but do not unambiguously explain the developed phage resistance.

Bacterial defence mechanisms against ssDNA phages are partly the same than those against dsDNA phages. Especially extracellular defences such as surface modifications, and phage receptor-presenting vesicles, but also some intracellular mechanisms like abortive infection, are used by bacteria to defend against both phage types (“Defense and anti-defense mechanisms of bacteria and bacteriophages,” 2024a; “Defense and anti-defense mechanisms of bacteria and bacteriophages,” 2024b; Gao and Feng, 2023; Sasaki et al., 2023). The mechanisms targeted directly against phage genomes such as restriction-modification systems and CRISPR-Cas systems most commonly cleave specifically dsDNA (Labrie et al., 2010). It is worth noting that even systems acting against dsDNA might potentially weaken the replication of ssDNA phages, including FLiP. The conserved replication initiation protein (Rep) encoding gene found in FLiP genome refers to the rolling circle replication (RCR) system (Laanto et al., 2017b), a common replication mechanism among ssDNA phages such as *Microviridae* and *Inoviridae* (Nguyen et al., 2023). In this replication system, the ssDNA genome is first transformed into replicative form (RF), which has a structure similar to a dsDNA plasmid, meaning that infected cells do also contain FLiP genome in dsDNA form. Thus, the presence of RFs may allow the dsDNA-targeting anti-phage systems to be used against ssDNA phages during DNA replication. On the other hand, these systems cannot destroy the newly formed ssDNA genomes, and consequently, they might not offer a complete defence against ssDNA phages.

In a normal lytic cycle RFs are short-lived, temporary structures, and the other strand will rapidly be nicked by Rep, displaced and circularized into new ssDNA genomes. Interestingly, the unintegrated FLiP genomes in dsDNA form are not always short lived. We showed the presence of FLiP as a circular extrachromosomal dsDNA element within the resistant B114r. This finding was further supported by the discovery of FLiP genome in the whole genome sequencing data (206x coverage). These results refer to a lysogenic state rather than to RFs, as lysogeny may provide the host with strong superinfection exclusion, which could explain the developed resistance. Because the population of progeny cells was permanently resistant through several generations (data not shown) the evidence of true lysogeny is stronger than to more transient states like RFs or pseudolysogeny. These findings align well with the facts that lysogeny is a stable and often long-term state, maintained by phage-encoded repressors, and can involve the phage genome either integrating into the host genome or existing as a plasmid-like extrachromosomal element (Howard-Varona et al., 2017). We did find a putative repressor gene in FLiP genome (Mäkelä et al., 2025; ORF04), pointing further into the direction that lysogeny is possible for this phage. In lysogeny, the prophage is replicated along with the host cell division, ensuring all daughter cells receive a copy which results in consistent resistance in the population. As a conclusion, lysogeny is probable for FLiP in B114 resistant variant (B114r), given that the resistance was provided by superinfection exclusion and not by activation of other resistance mechanisms.

Although sequencing analysis did not identify FLiP reads in the ancestral hosts, plasmid DNA gave a low signal in FLiP-specific PCR, suggesting that small percentage of the cells in the susceptible populations, which have never been exposed to FLiP in the laboratory, might contain FLiP in circular dsDNA form as well. This suggests that coexistence of some lysogens or pseudolysogens among the mainly susceptible host population is possible also in nature in case of FLiP-like phages. Nevertheless, the extrachromosomal presence of FLiP depends also on the host. Here, we studied three hosts (B330, B114 and B167), and only in one (B114) the PCR-signals of FLiP presence were strong. This indicates that in the other hosts (which are more closely related together), a smaller proportion carried FLiP. Furthermore, primers amplifying FLiP MCP produced positive results in PCR for seven other *Flavobacterium* strains, which had been isolated from environmental sources and fish farms. It is possible that FLiP-type temperate phages are more abundant among flavobacteria in various aquatic ecosystems than currently acknowledged (Yutin et al., 2018).

While lysogeny can be rather unambiguously defined, pseudolysogenic presence in an infected cell is more elusive (Cenens et al., 2013a; Mäntynen et al., 2021). Pseudolysogeny is an unstable state and does not commit to the lytic or lysogenic cycle until a change in the growth conditions occurs (Ripp&Miller 1997, Ripp&Miller 1998, Los et al 2003). The phage genome is transiently inactive, but not integrated into the host genome, and therefore unequally distributed among daughter cells during division, as observed e.g. in *Salmonella typhimurium* (Cenens et al., 2013b). Delayed lysis under stress and a rapid transition back to lytic cycle when conditions improve, were observed in FLiP-infections, and may be considered as indications of pseudolysogeny (Ripp and Miller, 1997; Wilson et al., 1996). This kind of phenomena could, on the other hand, be caused simply by slower replication under unfavorable conditions. The FLiP dsDNA plasmid-like structures in the infected cells may thus be phage genomes in RF form, whose typically brief presence may be prolonged under stressful conditions due to slower infection. Alternatively, they may be truly pseudolysogenic extrachromosomal forms of the phage, that is further distributed into proportion of daughter cells during cell division (Cenens et al 2013), thus allowing FLiP to survive in the population. Lysogeny is known to provide host cells with strong protection against infections by similar phages in some phage-bacterium pairs (Howard-Varona et al., 2017; Labrie et al., 2010). However, comparable research on pseudolysogeny in this context is largely lacking. Yet, it appears that significantly different phage genes are expressed during lysogeny and pseudolysogeny (Cenens et al., 2013a), suggesting the possible superinfection immunity provided by a pseudolysogenic phage might be achieved through different mechanisms compared to lysogeny. Future work should clarify whether FLiP can establish lysogeny or pseudolysogeny and how transitions between these states and lytic infection are regulated, for example by comparing expression of candidate genes such as a putative repressor across different infection contexts.

Although phages are often seen as parasites of bacteria, mutualistic relationships are also well known (Obeng et al., 2016). Both ssDNA and dsDNA phages can establish mutualistic relationships to their hosts as temperate phages (Sausset et al., 2020) providing the host with fitness advantages such as antibiotic resistance, toxin genes or mediating superinfection exclusion (Jian et al., 2021; McLeod et al., 2005). Although the possible benefits of FLiP as circular extrachromosomal dsDNA element in *Flavobacterium* populations in nature is not exactly known, our study adds another example on the complexity of phage-host interactions and phage life cycles. The dsDNA plasmid-like form residing within the host may be the key to the flexibility of FLiP-host-interactions, giving FLiP means to rapidly switch between lytic infection, lysogeny, and pseudolysogeny, enabling immediate response to changing conditions and effective competition with much larger phages.

## Acknowledgements

We want to thank Dr. Heidi Kunttu, Dr. Roghaieh Ashrafi, Dr. Päivi Rintamäki (University of Oulu), Eva Jansson (National Veterinary Institute of Sweden), Yrjö Lankinen, Natural research Institute Finland and Mark McBride (UWM, USA) for providing bacterial strains. We thank Petri Papponen for technical assistance in the laboratory. The research was funded by research grants from Emil Aaltonen Foundation (#200260 L.-R.S.), Research Council of Finland (#346772, L.-R.S. and #354982, R.P.), Olvi Foundation (#201910409 K.M.) and The Finnish Concordia Fund (#20200077 K.M.). This project has received funding from the European Research Council (ERC) under the European Union’s Horizon Europe research and innovation programme (grant agreement No 101117204) (E.L.).

