## Supporting information file 1 for "ssDNA phage FLiP resides in dsDNA form in resistant *Flavobacterium* host"

### PCR details:

**Primer sequences (5’-3’):**

| | |
|---|---|
| FLiP MCP Forward | GAATGTTGTTCGCGGTGCTT |
| FLiP MCP Reverse | CGACCAATGGGAAGAGGGAG |
| FLiP Rep Forward | TCAGCGCAAAGGTTAGGCAT |
| FLiP Rep Reverse | GCTGTGCTAACGCCCAAATC |

**PCR program (MCP and Rep):**

| | | |
|---|---|---|
| 95°C | 7 min | |
| 95°C | 30 s | 30x |
| 62°C | 30 s | 30x |
| 72°C | 1 min | 30x |
| 72°C | 10 min | |
| 12°C | ∞ | |

Table S1. Bacterial strains included in the PCR analysis using primers amplifying either FLiP major capsid protein (MCP) or replication initiation protein (Rep) gene.

| Flavobacterium strain | Species | Sampling site | Year | | MCP PCR positive | Rep PCR positive |
|---|---|---|---|---|---|---|
| UW101 | *F. johnsoniae* | United Kingdom | not known | (McBride et al., 2009) | | |
| B28 | sp. | Lake Konnevesi | 2006 | (Laanto et al., 2011) | | |
| B067 | *F. columnare* | Fish farm, Central Finland | 2007 | (Laanto et al., 2011) | X | |
| B108 | sp. | River Vantaanjoki | 2008 | unpubl | | |
| B121 | sp. | River Tsarsjoki | 2008 | (Laanto et al., 2011) | | |
| B127 | sp. | Lake Kevojärvi | 2008 | (Laanto et al., 2011) | | |
| B130 | sp. | Lake Kevojärvi | 2008 | (Laanto et al., 2011) | | |
| B138 | sp. | Small pond in Central Finland | 2008 | unpubl | X | |
| B169 | sp. | Lake Jyväsjärvi | 2008 | (Laanto et al., 2011) | | |
| B190 | sp. | Fish farm, Central Finland | 2008 | unpubl | | |
| B196 | sp. | Fish farm, Central Finland | 2008 | unpubl | X | |
| B207 | sp. | Lake Inari | 2009 | (Laanto et al., 2011) | X | |
| B222 | sp. | River Kymijoki | 2009 | (Laanto et al., 2011) | X | |
| B223 | sp. | River Kymijoki | 2009 | (Laanto et al., 2011) | | |
| B224 | sp. | River Kymijoki | 2009 | (Laanto et al., 2011) | | |
| FCO-F86 | *F. columnare* | Fish farm, North Finland | 2017 | (Runtuvuori-Salmela et al., 2022) | | |
| FCO-F93 | *F. columnare* | Fish farm, North Finland | 2017 | (Runtuvuori-Salmela et al., 2022) | | |
| FCO-F133 | *F. columnare* | Fish farm, North Finland | 2018 | unpubl | | |
| FCO-F134 | *F. columnare* | Fish farm, North Finland | 2018 | unpubl | | |
| FCO-F155 | *F. columnare* | Fish farm, Central Finland | 2019 | unpubl | | |
| B350 | *F. columnare* | Fish farm outlet water, Central Finland | 2010 | (Sundberg et al., 2016) | X | |
| B430 | *F. columnare* | Fish farm, Central Finland | 2003 | (Ashrafi et al., 2015) | | |
| B480 | *F. columnare* | Fish farm, Central Finland | 2012 | (Runtuvuori-Salmela et al., 2022) | X | |
| 3/3449 | *F. columnare* | Fish farm, Sweden | 2017 | (Runtuvuori-Salmela et al., 2022) | | |
| 4/3450 | *F. columnare* | Fish farm, Sweden | 2017 | (Runtuvuori-Salmela et al., 2022) | | |
| 573451 | *F. columnare* | Fish farm, Sweden | 2017 | (Runtuvuori-Salmela et al., 2022) | | |
| 5/3460 | *F. columnare* | Fish farm, Sweden | 2017 | (Runtuvuori-Salmela et al., 2022) | | |
| 6/3461 | *F. columnare* | Fish farm, Sweden | 2017 | (Runtuvuori-Salmela et al., 2022) | | |
| A2I | sp. | Fish farm, Central Finland | 2018 | unpubl | | |
| A6I | sp. | Fish farm, Central Finland | 2018 | unpubl | | |
| A8I | sp. | Fish farm, Central Finland | 2018 | unpubl | | |
| F | *F. columnare* | | | *F. columnare* type strain NCIMB 2248 | | |

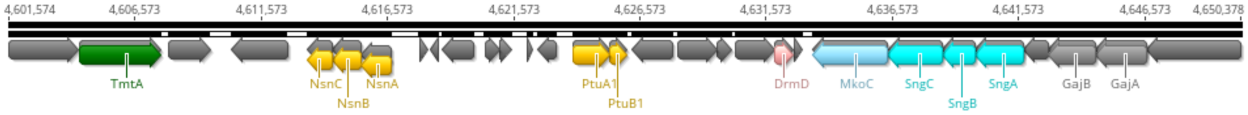


Figure S1. Defense island of *Flavobacterium* sp. B114r. The region includes defense systems Tiamat (genes coding for TmtA), Menshen (NsnA, NsnB, NsnC), Septu (PtuA1, PtuB1), DMS (DrmD), Mokosh (MkoC), Shango (SngA, SngB, SngC) and Gabija (GajA, GajB).

Ashrafi, R., Pulkkinen, K., Sundberg, L.-R., Pekkala, N., Ketola, T., 2015. A multilocus sequence analysis scheme for characterization of Flavobacterium columnare isolates. BMC Microbiology 15, 243. https://doi.org/10.1186/s12866-015-0576-4

Laanto, E., Sundberg, L.-R., Bamford, J.K.H., 2011. Phage Specificity of the Freshwater Fish Pathogen Flavobacterium columnare ▿. Appl Environ Microbiol 77, 7868–7872. https://doi.org/10.1128/AEM.05574-11

McBride, M.J., Xie, G., Martens, E.C., Lapidus, A., Henrissat, B., Rhodes, R.G., Goltsman, E., Wang, W., Xu, J., Hunnicutt, D.W., Staroscik, A.M., Hoover, T.R., Cheng, Y.-Q., Stein, J.L., 2009. Novel Features of the Polysaccharide-Digesting Gliding Bacterium Flavobacterium johnsoniae as Revealed by Genome Sequence Analysis. Applied and Environmental Microbiology 75, 6864–6875. https://doi.org/10.1128/AEM.01495-09

Runtuvuori-Salmela, A., Kunttu, H.M.T., Laanto, E., Almeida, G.M.F., Mäkelä, K., Middelboe, M., Sundberg, L., 2022. Prevalence of genetically similar Flavobacterium columnare phages across aquaculture environments reveals a strong potential for pathogen control. Environ Microbiol 24, 2404–2420. https://doi.org/10.1111/1462-2920.15901

Sundberg, L.-R., Ketola, T., Laanto, E., Kinnula, H., Bamford, J.K.H., Penttinen, R., Mappes, J., 2016. Intensive aquaculture selects for increased virulence and interference competition in bacteria. Proceedings of the Royal Society B: Biological Sciences 283, 20153069. https://doi.org/10.1098/rspb.2015.3069
