## Supplementary Table 1 for "ssDNA phage FLiP resides in dsDNA form in resistant *Flavobacterium* host"

**Supplementary Table 1. Detected genetic changes between ancestral (B114, B167, B330) and the respective resistant variants (B114r, B167r, B330r) using breseq.**

***B114 vs. B114r***

| Position | Mutation | Annotation | Description |
|---|---|---|---|
| 52,084 | $(A)_{10\rightarrow9}$ | intergenic (+143/+43) | transketolase Cterminal -domaincontaining- protein/hypothetical protein |
| 1,397,503 | $(A)_{10\rightarrow9}$ | intergenic (-175/+69) | NADHquinone oxidoreductase subunit A/cold shock -domaincontaining- protein |
| 2,487,895 | $(T)_{9\rightarrow8}$ | intergenic (-31/+305) | glycoside hydrolase family 2 TIM barreldomain containing protein/RagB/SusD- family nutrient uptake outer membrane protein |
| 2,488,892 | Δ1 bp | coding (28/720 nt) | RagB/SusD family nutrient uptake outer membrane protein |
| 2,489,001 | +A | coding (138/141 nt) | hypothetical protein |
| 2,489,010 | Δ1 bp | coding (129/141 nt) | hypothetical protein |
| 2,489,015 | +T | coding (124/141 nt) | hypothetical protein |
| 2,489,217 | +TT | coding (268/279 nt) | RagB/SusD family nutrient uptake outer membrane protein |
| 2,489,489 | +A | coding (205/219 nt) | hypothetical protein |
| 2,680,145 | $(T)_{10\rightarrow9}$ | intergenic (+76/-1280) | preprotein translocase subunit YajC/IS4 family transposase |
| 4,252,344 | $(T)_{9\rightarrow8}$ | intergenic (+73/+101) | META domaincontaining protein/-TonBdependent- receptor |
| 5,527,951 | $(T)_{10\rightarrow9}$ | intergenic (+67/-320) | hypothetical protein/M15 family metallopeptidase |

***B167 vs. B167r***

| Position | Mutation | Annotation | Description |
|---|---|---|---|
| 597,706 | (A)$_{8\rightarrow9}$ | intergenic (-125/-97) | tyrosine-tRNA ligase/-NADdependent- epimerase/dehydratase family protein |
| 1,146,451 | (T)$_{7\rightarrow8}$ | intergenic (+563/-79) | AAA family ATPase/GNAT family N-acetyltransferase |
| 2,319,336 | (T)$_{15\rightarrow16}$ | intergenic (+87/-541) | NUDIX hydrolase/TonBdependent- receptor |
| 2,902,723 | A→T | I153F (ATC→TTC) | cysteine synthase family protein |
| 3,558,383 | (T)$_{6\rightarrow7}$ | coding (4052/4098 nt) | twocomponent regulator propeller -domaincontaining- protein |
| 5,102,172 | +A | coding (419/495 nt) | DUF4369 domaincontaining- protein |
| 5,102,604 | +T | intergenic (-14/+537) | DUF4369 domaincontaining- protein/response regulator |
| 5,103,005 | (T)$_{6\rightarrow7}$ | intergenic (-415/+136) | DUF4369 domaincontaining- protein/response regulator |
| 5,103,427 | +C | intergenic (-26/+164) | response regulator/ATPbinding- protein |
| 5,103,572 | +A | intergenic (-171/+19) | response regulator/ATPbinding- protein |
| 5,103,978 | +T | coding (350/366 nt) | histidine kinase dimerization/phosphoacceptor -domaincontaining- protein |
| 5,104,444 | +G | coding (1818/1830 nt) | PAS domain Sbox- protein |
| 5,106,303 | +TA | intergenic (-42/+457) | PAS domain Sbox- protein/hypothetical protein |
| 5,106,639 | Δ1 bp | intergenic (-378/+121) | PAS domain Sbox- protein/hypothetical protein |
| 5,106,766 | T→C | K42K (AAA→AAG) | hypothetical protein |
| 5,106,852 | +GC | coding (40/132 nt) | hypothetical protein |
| 5,107,231 | +TAG | intergenic (-340/-41) | hypothetical protein/FADdependent- oxidoreductase |
| 5,109,009 | +T | coding (203/273 nt) | SDR family NAD(P)-dependent oxidoreductase |

***B330 vs. B330r***

| Position | Mutation | Annotation | Description |
|---|---|---|---|
| 66,382 | A→C | I408M (ATT→ATG) | AarF/UbiB family protein |
| 1,257,192 | T→C | R174G (AGA→GGA) | DUF3078 domaincontaining- protein |
| 2,753,321 | $(A)_{9\rightarrow10}$ | coding (394/399 nt) | curlilike amyloid fiber formation chaperone -CsgH |
| | | coding (15/354 nt) | CsgE family curlitype- amyloid fiber assembly protein |
| 2,883,551 | $(TAAAATC)_{23\rightarrow22}$ | intergenic (-279/-105) | ATPbinding cassette -domaincontaining- protein/hypothetical protein |
| 2,889,818 | T→C | E141G (GAA→GGA) | D-2-hydroxyacid dehydrogenase |
| 3,937,590 | $(A)_{5\rightarrow6}$ | pseudogene (521/1247 nt) | mechanosensitive ion channel |
| 4,913,045 | $(A)_{7\rightarrow8}$ | intergenic (-114/-5) | rhomboid family intramembrane serine protease/lysophospholipid acyltransferase family protein |
| 5,487,897 | $(A)_{6\rightarrow7}$ | coding (1421/1452 nt) | HD domaincontaining- protein |
