## Supplementary Table 2 for "ssDNA phage FLiP resides in dsDNA form in resistant *Flavobacterium* host"

| #Re | Host | seqid | system | target.name | hmm.accession | hmm.name | protein.name | full.seq.E.valu | domain.iE.valu | target.coverag | hmm.coverag | start | end | strand | target.descript | relative.positio | contig.end | all.domains | best.hits |
|---|---|---|---|---|---|---|---|---|---|---|---|---|---|---|---|---|---|---|---|
| | B114r | CP124858 | gabija | QLQ94_19350 | PDLC00070 | GajA_00010 | GajA | 1.5999999999 | 1.7999999999 | 0.982 | 0.997 | 4644605 | 4646618 | - | - | 3767 | 4712 | PDLC00061.GajA_00001, 1, 3.8e-06, 0.09, 0.09 \| PDLC00061.GajA_00001, 2, 0.0024, 0.196, 0.209 \| | PDLC00070, GajA_00010, 1.8e-189, 0.982, 0.997 \| PDLC00064, GajA_00004, 3.3e-48, 0.482, 0.726 \| PDLC00071, GajA_00011, 2.5e-40, 0.482, 0.537 \| PDLC00096, GajA_00036, 1.1e-39, 0.481, 0.525 \| PDLC00091, GajA_00031, 7.2e-38, 0.479, 0.525 |
| | B114r | CP124858 | gabija | QLQ94_19345 | PDLC00117 | GajB_00015 | GajB | 2,00E-186 | 2.2999999999 | 0.972 | 0.988 | 4642762 | 4644613 | - | - | 3766 | 4712 | PDLC00103.GajB_00001, 1, 3.3e-47, 0.917, 0.928 \| PDLC00104.GajB_00002, 1, 1.5e-23, 0.445, 0.408 | PDLC00117, GajB_00015, 2.3e-186, 0.972, 0.988 \| PDLC00127, GajB_00025, 6.1e-67, 0.911, 0.946 \| PDLC00130, GajB_00028, 5.4e-63, 0.898, 0.952 \| PDLC00123, GajB_00021, 8.2e-59, 0.938, 0.965 \| PDLC00129, GajB_00027, 9.3e-52, 0.898, 0.936 |
| | B114r | CP124858 | septu_type_I | QLQ94_19280 | PDLC00265 | PtuA1_00001 | PtuA1 | 8.6e-91 | 1.0999999999 | 0.99 | 0.977 | 4623916 | 4625398 | + | - | 3753 | 4712 | PDLC00265.PtuA1_00001, 1, 1.1e-90, 0.99, 0.977 \| PDLC00266.PtuA1_00002, 1, 1.8e-40, 0.959, 0.96 | PDLC00265, PtuA1_00001, 1.1e-90, 0.99, 0.977 \| PDLC00283, PtuA1_00019, 9e-47, 0.947, 0.94 \| PDLC00266, PtuA1_00002, 1.8e-40, 0.959, 0.961 \| PDLC00285, PtuA1_00021, 2.1e-33, 0.961, 0.963 \| PDLC00303, PtuA1_00039, 5.9e-29, 0.525, 0.635 |
| | B114r | CP124858 | septu_type_I | QLQ94_19285 | PDLC00325 | PtuB1_00014 | PtuB1 | 2.3999999999 | 2.9e-46 | 0.889 | 0.934 | 4625394 | 4626072 | + | - | 3754 | 4712 | PDLC00288.PtuA1_00024, 1, 4.6e-09, 0.609, 0.199 \| PDLC00292.PtuA1_00028, 1, 3.5e-11, 0.76, 0.2 | PDLC00325, PtuB1_00014, 2.9e-46, 0.889, 0.934 \| PDLC00345, PtuB1_00034, 3.6e-45, 0.791, 0.789 \| PDLC00335, PtuB1_00024, 3.4e-39, 0.84, 0.907 \| PDLC00350, PtuB1_00039, 4.3e-27, 0.604, 0.607 \| PDLC00354, PtuB1_00043, 2.4e-20, 0.684, 0.804 |
| | B114r | CP124858 | DRT_class_II | QLQ94_05835 | PDLC02173 | Drt2_00001 | Drt2 | 1.7999999999 | 2.2999999999 | 0.967 | 0.995 | 1309182 | 1310712 | - | - | 1126 | 4712 | PDLC02173.Drt2_00001, 1, 2.3e-75, 0.967, 0.995 \| PDLC03754.drt-UG17, 1, 720, 0.014, 0.016 \| PDL | PDLC03757, drt-UG2, 1.1e-112, 0.978, 0.793 \| PDLC02173, Drt2_00001, 2.3e-75, 0.967, 0.995 \| PDLC03766, drt-UG28, 2.6e-20, 0.528, 0.391 \| PDLC03754, drt-UG17, 4.2e-18, 0.361, 0.326 \| PDLC03764, drt-UG26, 3.2e-16, 0.373, 0.397 |
| | B114r | CP124858 | DMS_other | QLQ94_19310 | PDLC02630 | DrmD_00013 | DrmD | 1.8000000000 | 2.3000000000 | 0.955 | 0.885 | 4631915 | 4632644 | + | - | 3759 | 4712 | PDLC02630.DrmD_00013, 1, 2.3e-27, 0.955, 0.885 \| PDLC03192.Specificity_I_00141, 1, 0.015, 0.298 | PDLC02630, DrmD_00013, 2.3e-27, 0.955, 0.885 \| PDLC04140, mads_8, 1.9e-05, 0.384, 0.274 \| PDLC03192, Specificity_I_00141, 0.015, 0.298, 0.172 |
| | B114r | CP124858 | DRT_other | QLQ94_05840 | PDLC02926 | retron_type_VI_hth_cluster2_1 | HTH_VI | 8,00E-15 | 1,00E-14 | 0.689 | 0.674 | 1311302 | 1311575 | - | - | 1127 | 4712 | PDLC02926,retron_type_VI_hth_cluster2_1, 1, 1e-14, 0.689, 0.674 | PDLC02926, retron_type_VI_hth_cluster2_1, 1e-14, 0.689, 0.674 |
| | B114r | CP124858 | DRT_other | QLQ94_22500 | PDLC02926 | retron_type_VI_hth_cluster2_1 | HTH_VI | 5.7e-10 | 8.2e-10 | 0.474 | 0.696 | 5434234 | 5434636 | + | - | 4369 | 4712 | PDLC02926,retron_type_VI_hth_cluster2_1, 1, 8.2e-10, 0.474, 0.696 | PDLC02926, retron_type_VI_hth_cluster2_1, 8.2e-10, 0.474, 0.696 |
| | B114r | CP124858 | RM_type_I | QLQ94_03225 | PDLC03025 | MTase_I_00008 | MTase_I | 1.3999999999 | 5.0999999999 | 0.925 | 0.784 | 701216 | 702653 | + | - | 613 | 4712 | PDLC03018.MTase_I_00001, 1, 5.7e-13, 0.209, 0.131 \| PDLC03018.MTase_I_00001, 2, 6, 0.1, 0.061 \| | PDLC03025, MTase_I_00008, 5.1e-182, 0.925, 0.784 \| PDLC03021, MTase_I_00004, 1.4e-158, 0.99, 0.99 \| PDLC03026, MTase_I_00009, 1.1e-66, 0.941, 0.918 \| PDLC03036, MTase_I_00019, 4.7e-43, 0.797, 0.666 \| PDLC03022, MTase_I_00005, 8.4e-33, 0.732, 0.422 |
| | B114r | CP124858 | RM_type_I | QLQ94_03215 | PDLC03049 | REase_I_00010 | REase_I | 0 | 0 | 0.995 | 0.986 | 697613 | 700055 | + | - | 611 | 4712 | PDLC03042.REase_I_00003, 1, 0.23, 0.054, 0.038 \| PDLC03042.REase_I_00003, 2, 2.5e-37, 0.513, 0 | PDLC03049, REase_I_00010, 0, 0.995, 0.986 \| PDLC03050, REase_I_00011, 1.4e-65, 0.24, 0.89 \| PDLC03042, REase_I_00003, 2.5e-37, 0.513, 0.364 \| PDLC03047, REase_I_00008, 4.9e-28, 0.624, 0.442 \| PDLC03045, REase_I_00006, 5.1e-13, 0.269, 0.212 |
| | B114r | CP124858 | RM_type_I | QLQ94_03230 | PDLC03076 | Specificity_I_00025 | Specificity_I | 1,00E-46 | 2.0999999999 | 0.394 | 0.504 | 702649 | 704599 | + | - | 614 | 4712 | PDLC03038.MTase_I_00021, 1, 0.0026, 0.177, 0.125 \| PDLC03039.MTase_I_00022, 1, 2.5e-14, 0.274 | PDLC03056, Specificity_I_00005, 6.6e-62, 0.408, 0.447 \| PDLC03179, Specificity_I_00128, 4.7e-52, 0.408, 0.445 \| PDLC03076, Specificity_I_00025, 2.1e-40, 0.394, 0.504 \| PDLC03169, Specificity_I_00118, 3.9e-36, 0.612, 0.766 \| PDLC03096, Specificity_I_00045, 4.9e-30, 0.293, 0.284 |
| | B114r | CP124858 | RM_type_IV | QLQ94_06760 | PDLC03659 | mREase_IV_00005 | mREase_IV | 1.2999999999 | 1.5999999999 | 0.881 | 0.981 | 1521111 | 1524261 | - | - | 1306 | 4712 | PDLC03658.mREase_IV_00004, 1, 2.9e-82, 0.592, 0.644 \| PDLC03658.mREase_IV_00004, 2, 46, 0.1 | PDLC03659, mREase_IV_00005, 1.6e-231, 0.881, 0.981 \| PDLC03658, mREase_IV_00004, 2.9e-82, 0.592, 0.644 |
| | B114r | CP124858 | DRT_class_II | QLQ94_03205 | PDLC03778 | drt-UG39 | RT_UG39 | 1.0999999999 | 1.1999999999 | 0.993 | 0.993 | 693083 | 694850 | + | - | 609 | 4712 | PDLC02173.Drt2_00001, 1, 3.7e-13, 0.519, 0.685 \| PDLC02173.Drt2_00001, 2, 1400, 0.039, 0.054 \| P | PDLC03778, drt-UG39, 1.2e-170, 0.993, 0.993 \| PDLC03766, drt-UG28, 2.7e-24, 0.524, 0.496 \| PDLC03757, drt-UG2, 5e-22, 0.673, 0.627 \| PDLC03754, drt-UG17, 9e-14, 0.306, 0.312 \| PDLC02173, Drt2_00001, 3.7e-13, 0.519, 0.685 |
| | B114r | CP124858 | DRT class III | QLQ94_22510 | PDLC03780 | drt-UG5_large | RT_UG5-nitrilase | 3.3999999999 | 8.2999999999 | 0.991 | 0.964 | 5435753 | 5439071 | - | - | 4371 | 4712 | PDLC02995.RT, 1, 0.34, 0.07, 0.072 \| PDLC02995.RT, 2, 9.3e-25, 0.319, 0.371 \| PDLC02995.RT, 3, 6 | PDLC03780, drt-UG5_large, 8.3e-227, 0.991, 0.964 \| PDLC02995, RT, 6.8e-58, 0.258, 0.303 \| PDLC03781, drt-UG5_small, 2.9e-54, 0.274, 0.475 \| PDLC03804, drt-cluster285, 1.9e-31, 0.229, 0.589 \| PDLC03745, drt-UG1, 5.1e-12, 0.192, 0.181 |
| | B114r | CP124858 | DRT_class_II | QLQ94_03200 | PDLC03818 | drt-cluster64 | RTasVirE-PriCT2 | 1.4999999999 | 1.7999999999 | 0.909 | 0.988 | 691851 | 692910 | - | - | 608 | 4712 | PDLC03818,drt-cluster64, 1, 1.8e-97, 0.909, 0.988 | PDLC03818, drt-cluster64, 1.8e-97, 0.909, 0.988 |
| | B114r | CP124858 | Menshen | QLQ94_19240 | PDLC03877 | Menshen__NsnA | NsnA | 1.8999999999 | 9.2999999999 | 0.816 | 0.842 | 4615520 | 4616714 | - | - | 3745 | 4712 | PDLC03877.Menshen__NsnA, 1, 5.3, 0.068, 0.07 \| PDLC03877.Menshen__NsnA, 2, 9.3e-101, 0.816, | PDLC03877, Menshen__NsnA, 9.3e-101, 0.816, 0.842 |
| | B114r | CP124858 | Menshen | QLQ94_19235 | PDLC03878 | Menshen__NsnB | NsnB | 1.5999999999 | 1.7999999999 | 0.995 | 0.991 | 4614406 | 4615543 | - | - | 3744 | 4712 | PDLC03878,Menshen__NsnB, 1, 1.8e-109, 0.995, 0.991 | PDLC03878, Menshen__NsnB, 1.8e-109, 0.995, 0.991 |
| | B114r | CP124858 | Menshen | QLQ94_19230 | PDLC03890 | Menshen__NsnC_2598643479 | NsnC | 5.2e-13 | 8.3e-13 | 0.332 | 0.326 | 4613417 | 4614377 | - | - | 3743 | 4712 | PDLC03890.Menshen__NsnC_2598643479, 1, 45, 0.254, 0.258 \| PDLC03890.Menshen__NsnC_25986 | PDLC03890, Menshen__NsnC_2598643479, 8.3e-13, 0.332, 0.326 \| PDLC03905, Menshen__NsnC_2656976156, 1.7e-10, 0.245, 0.264 \| PDLC03912, Menshen__NsnC_2705968471, 0.0059, 0.473, 0.465 |
| | B114r | CP124858 | Mokosh_Type | QLQ94_19320 | PDLC03924 | Mokosh_TypeII__MkoC | MkoC | 7.9999999999 | 8.9999999999 | 0.992 | 0.844 | 4633407 | 4636395 | - | - | 3761 | 4712 | PDLC03924,Mokosh_TypeII__MkoC, 1, 9e-293, 0.992, 0.844 | PDLC03924, Mokosh_TypeII__MkoC, 9e-293, 0.992, 0.844 |
| | B114r | CP124858 | Shango | QLQ94_19335 | PDLC03958 | Shango__SngA | SngA | 4.2e-44 | 7.4e-44 | 0.904 | 0.867 | 4639846 | 4641778 | - | - | 3764 | 4712 | PDLC03958,Shango__SngA, 1, 7.4e-44, 0.904, 0.867 | PDLC03958, Shango__SngA, 7.4e-44, 0.904, 0.867 |
| | B114r | CP124858 | Shango | QLQ94_19330 | PDLC03959 | Shango__SngB | SngB | 2.5999999999 | 3.0999999999 | 0.959 | 0.952 | 4638533 | 4639850 | - | - | 3763 | 4712 | PDLC03959,Shango__SngB, 1, 3.1e-164, 0.959, 0.952 | PDLC03959, Shango__SngB, 3.1e-164, 0.959, 0.952 |
| | B114r | CP124858 | Shango | QLQ94_19325 | PDLC03960 | Shango__SngC | SngC | 1.7999999999 | 2.2999999999 | 0.982 | 0.989 | 4636401 | 4638534 | - | - | 3762 | 4712 | PDLC03960,Shango__SngC, 1, 2.3e-210, 0.982, 0.989 | PDLC03960, Shango__SngC, 2.3e-210, 0.982, 0.989 |
| | B114r | CP124858 | SoFic | QLQ94_14145 | PDLC03963 | SoFic__SoFic | SoFic | 2.6e-32 | 4.3e-32 | 0.693 | 0.687 | 3312727 | 3313777 | - | - | 2741 | 4712 | PDLC03963,SoFic__SoFic, 1, 4.3e-32, 0.693, 0.687 | PDLC03963, SoFic__SoFic, 4.3e-32, 0.693, 0.687 |
| | B114r | CP124858 | SoFic | QLQ94_20895 | PDLC03963 | SoFic__SoFic | SoFic | 1,00E-28 | 1.1999999999 | 0.882 | 0.762 | 5026727 | 5027645 | + | - | 4067 | 4712 | PDLC03963,SoFic__SoFic, 1, 1.2e-28, 0.882, 0.762 | PDLC03963, SoFic__SoFic, 1.2e-28, 0.882, 0.762 |
| | B114r | CP124858 | Tiamat | QLQ94_19215 | PDLC03972 | Tiamat__TmtA_2752664533 | TmtA | 1.6999999999 | 4.2999999999 | 0.714 | 0.598 | 4604350 | 4607584 | + | - | 3740 | 4712 | PDLC03964.Tiamat__TmtA, 1, 3.2e-31, 0.731, 0.527 \| PDLC03964.Tiamat__TmtA, 2, 6.5e-15, 0.197, 0 | PDLC03972, Tiamat__TmtA_2752664533, 4.3e-56, 0.714, 0.598 \| PDLC03968, Tiamat__TmtA_2660206627, 1e-42, 0.331, 0.244 \| PDLC03966, Tiamat__TmtA_2599863134, 1.7e-32, 0.457, 0.321 \| PDLC03964, Tiamat__TmtA, 3.2e-31, 0.731, 0.527 \| PDLC03965, Tiamat__TmtA_2505168307, 2.6e-18, 0.075, 0.06 |
| | B114r | CP124858 | PDC-M62 | QLQ94_03235 | PDLC04735 | PDC-M62A_WP_217027783.1 | PDC-M62A | 2.6e-91 | 3.3e-91 | 0.991 | 0.995 | 704649 | 706617 | + | - | 615 | 4712 | PDLC04732.PDC-M62A_WP_092333670.1, 1, 2.5e-24, 0.971, 0.963 \| PDLC04733.PDC-M62A_WP_1 | PDLC04735, PDC-M62A_WP_217027783.1, 3.3e-91, 0.991, 0.995 \| PDLC04734, PDC-M62A_WP_001166610.1, 3.2e-63, 0.983, 0.978 \| PDLC04737, PDC-M62A_WP_152058451.1, 7.8e-55, 0.983, 0.983 \| PDLC04738, PDC-M62A_WP_088115436.1, 1.7e-29, 0.299, 0.263 \| PDLC04736, PDC-M62A_WP_051624115.1, 9.4e-27, 0.366, 0.346 |
| | B114r | CP124858 | PDC-M62 | QLQ94_03240 | PDLC04743 | PDC-M62B_WP_051775087.1 | PDC-M62B | 1.5e-59 | 1.7e-59 | 0.985 | 0.988 | 706594 | 707602 | + | - | 616 | 4712 | PDLC04740.PDC-M62B_WP_160541642.1, 1, 2100, 0.051, 0.044 \| PDLC04740.PDC-M62B_WP_160 | PDLC04743, PDC-M62B_WP_051775087.1, 1.7e-59, 0.985, 0.988 \| PDLC04741, PDC-M62B_WP_088115435.1, 4.7e-34, 0.696, 0.626 \| PDLC04742, PDC-M62B_WP_098312182.1, 7.2e-31, 0.985, 0.985 \| PDLC04740, PDC-M62B_WP_160541642.1, 3.5e-18, 0.618, 0.522 |
| | B114r | CP124858 | PDC-S02 | QLQ94_10840 | PDLC04795 | PDC-S02_WP_011937941.1 | PDC-S02 | 7.1999999999 | 7.8999999999 | 0.976 | 0.899 | 2548920 | 2549907 | - | - | 2087 | 4712 | PDLC04793.PDC-S02_WP_149946094.1, 1, 0.00011, 0.232, 0.667 \| PDLC04794.PDC-S02_WP_0912 | PDLC04795, PDC-S02_WP_011937941.1, 7.9e-131, 0.976, 0.899 \| PDLC04831, PDC-S02_WP_143549780.1, 7.1e-121, 0.957, 0.939 \| PDLC04822, PDC-S02_WP_207168644.1, 3.1e-100, 0.869, 0.959 \| PDLC04810, PDC-S02_WP_080642564.1, 3.5e-99, 0.924, 0.893 \| PDLC04803, PDC-S02_WP_226889553.1, 1.6e-93, 0.78, 0.941 |
| | B114r | CP124858 | PDC-S02 | QLQ94_00735 | PDLC04807 | PDC-S02_WP_151674975.1 | PDC-S02 | 8.0999999999 | 8.9999999999 | 0.988 | 0.974 | 150269 | 151310 | + | - | 136 | 4712 | PDLC04793.PDC-S02_WP_149946094.1, 1, 1.2e-17, 0.22, 0.658 \| PDLC04794.PDC-S02_WP_09125 | PDLC04807, PDC-S02_WP_151674975.1, 9e-129, 0.988, 0.974 \| PDLC04822, PDC-S02_WP_207168644.1, 1.8e-111, 0.873, 0.993 \| PDLC04831, PDC-S02_WP_143549780.1, 1.7e-89, 0.934, 0.925 \| PDLC04795, PDC-S02_WP_011937941.1, 3.3e-88, 0.98, 0.93 \| PDLC04803, PDC-S02_WP_226889553.1, 1.3e-81, 0.772, 0.959 |
| | B114r | CP124858 | PDC-S05 | QLQ94_06405 | PDLC04852 | PDC-S05_WP_110807286.1 | PDC-S05 | 1.5999999999 | 1.7999999999 | 0.988 | 0.962 | 1442485 | 1443490 | + | - | 1235 | 4712 | PDLC04843.PDC-S05_WP_181573034.1, 1, 5e-105, 0.985, 0.982 \| PDLC04844.PDC-S05_WP_0085 | PDLC04852, PDC-S05_WP_110807286.1, 1.8e-145, 0.988, 0.962 \| PDLC04853, PDC-S05_WP_228444512.1, 2e-132, 0.988, 0.982 \| PDLC04851, PDC-S05_WP_020395932.1, 8.5e-123, 0.964, 0.959 \| PDLC04868, PDC-S05_WP_205747975.1, 4.5e-108, 0.913, 0.934 \| PDLC04843, PDC-S05_WP_181573034.1, 5e-105, 0.985, 0.982 |
| | B114r | CP124858 | PDC-S05 | QLQ94_05165 | PDLC04856 | PDC-S05_WP_013135363.1 | PDC-S05 | 8.3e-71 | 9.0999999999 | 0.988 | 0.989 | 1162463 | 1162985 | - | - | 995 | 4712 | PDLC04849.PDC-S05_WP_217327273.1, 1, 6.6e-39, 0.913, 0.863 \| PDLC04856.PDC-S05_WP_0131 | PDLC04856, PDC-S05_WP_013135363.1, 9.1e-71, 0.988, 0.989 \| PDLC04869, PDC-S05_WP_078711833.1, 6.6e-55, 0.902, 0.529 \| PDLC04849, PDC-S05_WP_217327273.1, 6.6e-39, 0.913, 0.863 |
| | B114r | CP124858 | PDC-S07 | QLQ94_06765 | PDLC04899 | PDC-S07_WP_110056698.1 | PDC-S07 | 2.1999999999 | 2.3999999999 | 0.969 | 0.919 | 1524257 | 1524644 | - | - | 1307 | 4712 | PDLC04899,PDC-S07_WP_110056698.1, 1, 2.4e-48, 0.969, 0.919 | PDLC04899, PDC-S07_WP_110056698.1, 2.4e-48, 0.969, 0.919 |
| | B114r | CP124858 | PDC-S09 | QLQ94_19015 | PDLC04905 | PDC-S09_WP_017178955.1 | PDC-S09 | 1.2e-26 | 1.3e-26 | 0.96 | 0.941 | 4550682 | 4550982 | + | - | 3700 | 4712 | PDLC04902.PDC-S09_WP_203234077.1, 1, 3.5e-19, 0.919, 0.907 \| PDLC04903.PDC-S09_WP_0915 | PDLC04905, PDC-S09_WP_017178955.1, 1.3e-26, 0.96, 0.941 \| PDLC04906, PDC-S09_WP_015419311.1, 1.6e-20, 0.848, 0.827 \| PDLC04902, PDC-S09_WP_203234077.1, 3.5e-19, 0.919, 0.907 \| PDLC04904, PDC-S09_WP_046661845.1, 3.6e-17, 0.808, 0.74 \| PDLC04903, PDC-S09_WP_091549517.1, 9.3e-05, 0.323, 0.315 |
| | B114r | CP124858 | PDC-S15 | QLQ94_19205 | PDLC04932 | PDC-S15_WP_133278023.1 | PDC-S15 | 2.7999999999 | 3.9999999999 | 0.776 | 0.938 | 4600307 | 4601567 | + | - | 3738 | 4712 | PDLC04926.PDC-S15_WP_154556443.1, 1, 1.6e-06, 0.253, 0.317 \| PDLC04927.PDC-S15_WP_1136 | PDLC04932, PDC-S15_WP_133278023.1, 4e-66, 0.776, 0.938 \| PDLC04937, PDC-S15_WP_009831833.1, 9.9e-20, 0.411, 0.642 \| PDLC04931, PDC-S15_WP_226691295.1, 1.2e-19, 0.363, 0.514 \| PDLC04938, PDC-S15_WP_206299025.1, 8e-17, 0.346, 0.457 \| PDLC04940, PDC-S15_WP_171589529.1, 7.8e-15, 0.348, 0.5 |
| | B114r | CP124858 | PDC-S22 | QLQ94_01255 | PDLC04962 | PDC-S22_WP_003530377.1 | PDC-S22 | 1.2999999999 | 1.3999999999 | 0.989 | 0.954 | 261804 | 262620 | - | - | 236 | 4712 | PDLC04962,PDC-S22_WP_003530377.1, 1, 1.4e-132, 0.989, 0.954 | PDLC04962, PDC-S22_WP_003530377.1, 1.4e-132, 0.989, 0.954 |
| | B114r | CP124858 | PDC-S26 | QLQ94_03220 | PDLC04972 | PDC-S26_WP_074931224.1 | PDC-S26 | 8.0999999999 | 9.6999999999 | 0.972 | 0.982 | 700054 | 701224 | + | - | 612 | 4712 | PDLC04969.PDC-S26_WP_188767920.1, 1, 5e-07, 0.481, 0.493 \| PDLC04969.PDC-S26_WP_18876 | PDLC04972, PDC-S26_WP_074931224.1, 9.7e-99, 0.972, 0.982 \| PDLC04977, PDC-S26_WP_019505552.1, 2.6e-79, 0.956, 0.968 \| PDLC04971, PDC-S26_WP_155889373.1, 2.1e-49, 0.902, 0.894 \| PDLC04983, PDC-S26_WP_111444287.1, 2.2e-24, 0.871, 0.896 \| PDLC04980, PDC-S26_WP_095705398.1, 1.1e-22, 0.884, 0.913 |
| | B167r | CP124859 | retron_XIII | NA | - | TypeXIIIB_Ne144 | msr-msd | 4.9e-18 | NA | NA | NA | 5005873 | 5006100 | + | msr-msd ident | 4272.011 | 4613 | NA | NA |
| | B167r | CP124859 | AVAST type | QLQ96_10595 | PDLC02169 | Avs4_00001 | Avs4 | 8.1999999999 | 2.1e-117 | 0.836 | 0.878 | 2399080 | 2403943 | + | - | 2062 | 4613 | PDLC02169,Avs4_00001, 1, 2.1e-117, 0.836, 0.878 | PDLC02169, Avs4_00001, 2.1e-117, 0.836, 0.878 |
| | B167r | CP124859 | retron_XIII | QLQ96_21840 | PDLC02956 | retron_XIII | RT_XIII | 8.9999999999 | 9.8999999999 | 0.988 | 0.966 | 5006141 | 5007656 | + | - | 4273 | 4613 | PDLC02956,retron_XIII, 1, 9.9e-182, 0.988, 0.966 | PDLC02956, retron_XIII, 9.9e-182, 0.988, 0.966 |
| | B167r | CP124859 | retron_XIII | QLQ96_21850 | PDLC02972 | retron_type_XIII_znfinger_cluster114_1 | SWIM_XIII | 2.1999999999 | 2.7999999999 | 0.991 | 0.99 | 5010965 | 5012612 | + | - | 4275 | 4613 | PDLC02972,retron_type_XIII_znfinger_cluster114_1, 1, 2.8e-229, 0.991, 0.99 | PDLC02972, retron_type_XIII_znfinger_cluster114_1, 2.8e-229, 0.991, 0.99 |
| | B167r | CP124859 | retron_XIII | QLQ96_21845 | PDLC02973 | retron_type_XIII_wgr-heat_cluster216_1 | WGR_XIII | 0 | 0 | 0.998 | 0.994 | 5007674 | 5010962 | + | - | 4274 | 4613 | PDLC02920.retron_type_XIII_heat-arm_cluster216_2, 1, 1.8e-33, 0.993, 0.989 \| PDLC02922.retron_ty | PDLC02973, retron_type_XIII_wgr-heat_cluster216_1, 0, 0.998, 0.994 \| PDLC02920, retron_type_XIII_heat-arm_cluster216_2, 1.8e-33, 0.993, 0.989 \| PDLC02922, retron_type_XIII_heat-arm_cluster3262_1, 0.29, 0.157, 0.301 |
| | B167r | CP124859 | VSPR | QLQ96_03240 | PDLC03272 | MTase_II_00058 | MTase_II | 1.2999999999 | 1.3999999999 | 0.982 | 0.803 | 669635 | 670622 | + | - | 622 | 4613 | PDLC03217.MTase_II_00003, 1, 2.2e-57, 0.939, 0.447 \| PDLC03220.MTase_II_00006, 1, 1.3e-20, 0.4 | PDLC03272, MTase_II_00058, 1.4e-113, 0.982, 0.803 \| PDLC03378, MTase_II_00164, 1.3e-69, 0.976, 0.98 \| PDLC03361, MTase_II_00147, 1.6e-63, 0.976, 0.793 \| PDLC03217, MTase_II_00003, 2.2e-57, 0.939, 0.447 \| PDLC03234, MTase_II_00020, 6.5e-45, 0.57, 0.507 |
| | B167r | CP124859 | VSPR | QLQ96_03245 | PDLC03397 | NEase_II_00006 | Vsr | 1.4e-11 | 1.7e-11 | 0.819 | 0.66 | 670618 | 671002 | - | - | 623 | 4613 | PDLC03397,NEase_II_00006, 1, 1.7e-11, 0.819, 0.66 | PDLC03397, NEase_II_00006, 1.7e-11, 0.819, 0.66 |
| | B167r | CP124859 | DMS_other | QLQ96_12240 | PDLC03398 | NEase_II_00007 | REase_II | 9.1e-16 | 1.2e-15 | 0.604 | 0.606 | 2761615 | 2762284 | - | - | 2390 | 4613 | PDLC03398,NEase_II_00007, 1, 1.2e-15, 0.604, 0.606 | PDLC03398, NEase_II_00007, 1.2e-15, 0.604, 0.606 |
| | B167r | CP124859 | Lamassu Fan | QLQ96_10475 | PDLC03848 | Lamassu-Fam__LmuA_effector_Cap4_nuclease_II | LmuA | 1.6e-33 | 1.9e-33 | 0.939 | 0.938 | 2364616 | 2365741 | + | - | 2039 | 4613 | PDLC03848,Lamassu-Fam__LmuA_effector_Cap4_nuclease_II, 1, 1.9e-33, 0.939, 0.938 | PDLC03848, Lamassu-Fam__LmuA_effector_Cap4_nuclease_II, 1.9e-33, 0.939, 0.938 |
| | B167r | CP124859 | Lamassu Fan | QLQ96_10485 | PDLC03859 | Lamassu-Fam__LmuB_SMC_Cap4_nuclease_II | LmuB | 4.1999999999 | 4.6999999999 | 0.998 | 0.987 | 2366209 | 2368180 | + | - | 2041 | 4613 | PDLC00256.LmuB1_00008, 1, 2.1, 0.616, 0.707 \| PDLC00256.LmuB1_00008, 2, 2.7, 0.146, 0.167 \| PD | PDLC03859, Lamassu-Fam__LmuB_SMC_Cap4_nuclease_II, 4.7e-101, 0.998, 0.987 \| PDLC03863, Lamassu-Fam__LmuB_SMC_Mrr, 1.1e-28, 0.968, 0.951 \| PDLC03862, Lamassu-Fam__LmuB_SMC_Lipase, 5.7e-17, 0.966, 0.947 \| PDLC03867, Lamassu-Fam__LmuB_SMC_hypothetical, 4e-14, 0.619, 0.638 \| PDLC03866, Lamassu-Fam__LmuB_SMC_Sir2, 3.5e-11, 0.974, 0.956 |
| | B167r | CP124859 | Olokun | QLQ96_19790 | PDLC03925 | Olokun__OloA | OloA | 2.6e-56 | 3,00E-56 | 0.909 | 0.901 | 4520362 | 4522303 | + | - | 3877 | 4613 | PDLC03925,Olokun__OloA, 1, 3e-56, 0.909, 0.901 | PDLC03925, Olokun__OloA, 3e-56, 0.909, 0.901 |
| | B167r | CP124859 | Olokun | QLQ96_19795 | PDLC03926 | Olokun__OloB | OloB | 1.0999999999 | 1.1999999999 | 0.989 | 0.995 | 4522295 | 4524005 | + | - | 3878 | 4613 | PDLC03926,Olokun__OloB, 1, 1.2e-124, 0.989, 0.995 | PDLC03926, Olokun__OloB, 1.2e-124, 0.989, 0.995 |
| | B167r | CP124859 | SoFic | QLQ96_20750 | PDLC03963 | SoFic__SoFic | SoFic | 1.5e-31 | 1.8999999999 | 0.763 | 0.676 | 4759353 | 4760307 | + | - | 4065 | 4613 | PDLC03963,SoFic__SoFic, 1, 1.9e-31, 0.763, 0.676 | PDLC03963, SoFic__SoFic, 1.9e-31, 0.763, 0.676 |
| | B167r | CP124859 | DMS_other | QLQ96_12220 | PDLC04135 | mads_3 | mad3 | 1.0999999999 | 1.9e-41 | 0.344 | 0.968 | 2755786 | 2758582 | - | - | 2386 | 4613 | PDLC04135.mads_3, 1, 8.6e-13, 0.098, 0.267 \| PDLC04135.mads_3, 2, 1.5e-26, 0.223, 0.572 \| PDLC | PDLC04135, mads_3, 1.9e-41, 0.344, 0.968 \| PDLC04140, mads_8, 1.4e-12, 0.302, 0.732 |
| | B167r | CP124859 | PDC-M19 | QLQ96_04275 | PDLC04505 | PDC-M19A_WP_066957992.1 | PDC-M19A | 9.9999999999 | 1.0999999999 | 0.997 | 0.997 | 921589 | 922477 | + | - | 828 | 4613 | PDLC04504.PDC-M19A_WP_108422326.1, 1, 1.9e-100, 0.786, 0.991 \| PDLC04505.PDC-M19A_WP | PDLC04505, PDC-M19A_WP_066957992.1, 1.1e-125, 0.997, 0.997 \| PDLC04504, PDC-M19A_WP_108422326.1, 1.9e-100, 0.786, 0.991 |
| | B167r | CP124859 | PDC-M19 | QLQ96_04280 | PDLC04508 | PDC-M19B_WP_221312409.1 | PDC-M19B | 4.5999999999 | 5.1999999999 | 0.993 | 0.996 | 922479 | 923298 | + | - | 829 | 4613 | PDLC04506.PDC-M19B_WP_218031057.1, 1, 2.9e-50, 0.739, 0.966 \| PDLC04507.PDC-M19B_WP_0 | PDLC04508, PDC-M19B_WP_221312409.1, 5.2e-116, 0.993, 0.996 \| PDLC04506, PDC-M19B_WP_218031057.1, 2.9e-50, 0.739, 0.966 \| PDLC04507, PDC-M19B_WP_048156281.1, 1.2e-20, 0.974, 0.96 |
| | B167r | CP124859 | PDC-S05 | QLQ96_06235 | PDLC04852 | PDC-S05_WP_110807286.1 | PDC-S05 | 6.8999999999 | 7.5999999999 | 0.991 | 0.965 | 1369176 | 1370181 | + | - | 1215 | 4613 | PDLC04843.PDC-S05_WP_181573034.1, 1, 1.4e-106, 0.988, 0.985 \| PDLC04844.PDC-S05_WP_008 | PDLC04852, PDC-S05_WP_110807286.1, 7.6e-145, 0.991, 0.965 \| PDLC04853, PDC-S05_WP_228444512.1, 3.7e-132, 0.988, 0.982 \| PDLC04851, PDC-S05_WP_020395932.1, 1.3e-121, 0.964, 0.959 \| PDLC04868, PDC-S05_WP_205747975.1, 2.3e-107, 0.919, 0.94 \| PDLC04843, PDC-S05_WP_181573034.1, 1.4e-106, 0.988, 0.985 |
| | B167r | CP124859 | PDC-S05 | QLQ96_05110 | PDLC04856 | PDC-S05_WP_013135363.1 | PDC-S05 | 7,00E-70 | 7.6999999999 | 0.989 | 0.989 | 1118475 | 1119003 | - | - | 994 | 4613 | PDLC04849.PDC-S05_WP_217327273.1, 1, 4.7e-38, 0.92, 0.869 \| PDLC04856.PDC-S05_WP_01313 | PDLC04856, PDC-S05_WP_013135363.1, 7.7e-70, 0.989, 0.989 \| PDLC04869, PDC-S05_WP_078711833.1, 2.8e-55, 0.903, 0.529 \| PDLC04849, PDC-S05_WP_217327273.1, 4.7e-38, 0.92, 0.869 |
| | B167r | CP124859 | PDC-S13 | QLQ96_13840 | PDLC04919 | PDC-S13_WP_186842295.1 | PDC-S13 | 9.3999999999 | 9.9999999999 | 0.976 | 0.979 | 3083791 | 3084664 | + | - | 2708 | 4613 | PDLC04918.PDC-S13_WP_217993064.1, 1, 9.5e-06, 0.203, 0.276 \| PDLC04918.PDC-S13_WP_2179 | PDLC04919, PDC-S13_WP_186842295.1, 1e-87, 0.976, 0.979 \| PDLC04918, PDC-S13_WP_217993064.1, 1.3e-36, 0.41, 0.556 \| PDLC04920, PDC-S13_WP_097300085.1, 5.4e-10, 0.355, 0.364 |
| | B167r | CP124859 | PDC-S13 | QLQ96_05025 | PDLC04920 | PDC-S13_WP_097300085.1 | PDC-S13 | 1.3999999999 | 1.4999999999 | 0.978 | 0.93 | 1099220 | 1100042 | + | - | 977 | 4613 | PDLC04919.PDC-S13_WP_186842295.1, 1, 2.2e-05, 0.381, 0.374 \| PDLC04920.PDC-S13_WP_0973 | PDLC04920, PDC-S13_WP_097300085.1, 1.5e-121, 0.978, 0.93 \| PDLC04919, PDC-S13_WP_186842295.1, 2.2e-05, 0.381, 0.374 |
| | B330r | CP124860 | septu_type_I | QLQ95_05510 | PDLC00274 | PtuA1_00010 | PtuA1 | 2.0999999999 | 2.3999999999 | 0.998 | 0.998 | 424903 | 426184 | + | - | 411 | 4821 | PDLC00265.PtuA1_00001, 1, 0.0018, 0.31, 0.263 \| PDLC00265.PtuA1_00001, 2, 2.4e-16, 0.394, 0.34 | PDLC00274, PtuA1_00010, 2.4e-157, 0.998, 0.998 \| PDLC00289, PtuA1_00025, 3e-32, 0.974, 0.928 \| PDLC00271, PtuA1_00007, 1.5e-16, 0.981, 0.945 \| PDLC00265, PtuA1_00001, 2.4e-16, 0.394, 0.349 \| PDLC00283, PtuA1_00019, 3.3e-16, 0.272, 0.261 \| PDLC00290, PtuA1_00026, 3.3e-16, 0.838, 0.802 |
| | B330r | CP124860 | septu_type_I | QLQ95_05515 | PDLC00347 | PtuB1_00036 | PtuB1 | 9.6999999999 | 1.0999999999 | 0.984 | 0.99 | 426186 | 426765 | + | - | 412 | 4821 | PDLC00288.PtuA1_00024, 1, 0.15, 0.198, 0.052 \| PDLC00288.PtuA1_00024, 2, 0.08, 0.38, 0.101 \| PD | PDLC00347, PtuB1_00036, 1.1e-80, 0.984, 0.99 \| PDLC00332, PtuB1_00021, 1.8e-15, 0.802, 0.783 \| PDLC00337, PtuB1_00026, 7.7e-14, 0.88, 0.755 \| PDLC00346, PtuB1_00035, 3.4e-13, 0.74, 0.729 \| PDLC00322, PtuB1_00011, 2.6e-12, 0.781, 0.74 |
| | B330r | CP124860 | cbass type II | QLQ95_17890 | PDLC00718 | Effector_00056 | Effector | 2.7999999999 | 3.0999999999 | 0.986 | 0.986 | 3233811 | 3234687 | - | - | 2831 | 4821 | PDLC00715.Effector_00053, 1, 7.4e-63, 0.979, 0.952 \| PDLC00716.Effector_00054, 1, 9.8e-50, 0.973 | PDLC00718, Effector_00056, 3.1e-131, 0.986, 0.986 \| PDLC00715, Effector_00053, 7.4e-63, 0.979, 0.952 \| PDLC00721, Effector_00059, 1.9e-62, 0.976, 0.934 \| PDLC00716, Effector_00054, 9.8e-50, 0.973, 0.944 \| PDLC00723, Effector_00061, 2e-43, 0.962, 0.937 |
| | B330r | CP124860 | cbass type II | QLQ95_17905 | PDLC00736 | Effector_00074 | Effector | 2.3e-30 | 4.3e-30 | 0.906 | 0.531 | 3236560 | 3237652 | + | - | 2834 | 4821 | PDLC00736,Effector_00074, 1, 4.3e-30, 0.906, 0.531 | PDLC00736, Effector_00074, 4.3e-30, 0.906, 0.531 |
| | B330r | CP124860 | cbass type II | QLQ95_17920 | PDLC00736 | Effector_00074 | Effector | 2.8e-9 | 7.4e-9 | 0.331 | 0.351 | 3241963 | 3243805 | - | - | 2837 | 4821 | PDLC00736,Effector_00074, 1, 7.4e-09, 0.331, 0.351 | PDLC00736, Effector_00074, 7.4e-09, 0.331, 0.351 |
| | B330r | CP124860 | cbass type II | QLQ95_17940 | PDLC00752 | Effector_00090 | Effector | 1.4999999999 | 1.5999999999 | 0.998 | 0.996 | 3247420 | 3248950 | + | - | 2840 | 4821 | PDLC00676.Effector_00014, 1, 0.02, 0.216, 0.291 \| PDLC00688.Effector_00026, 1, 0.0015, 0.308, 0.4 | PDLC00752, Effector_00090, 1.6e-200, 0.998, 0.996 \| PDLC00737, Effector_00075, 8e-143, 0.998, 0.998 \| PDLC00735, Effector_00073, 1.4e-34, 0.481, 0.53 \| PDLC00741, Effector_00079, 4.1e-13, 0.358, 0.356 \| PDLC00744, Effector_00082, 4.2e-12, 0.536, 0.549 |
| | B330r | CP124860 | cbass type II | QLQ95_17895 | PDLC00802 | Cyclase_00042 | Cyclase | 1.3999999999 | 1.4999999999 | 0.982 | 0.994 | 3234683 | 3235691 | - | - | 2832 | 4821 | PDLC00769.Cyclase_00009, 1, 0.034, 0.257, 0.318 \| PDLC00769.Cyclase_00009, 2, 5.4e-12, 0.337, | PDLC00802, Cyclase_00042, 1.5e-139, 0.982, 0.994 \| PDLC00823, Cyclase_00063, 9.4e-47, 0.982, 0.993 \| PDLC00776, Cyclase_00016, 7.2e-22, 0.579, 0.666 \| PDLC00805, Cyclase_00045, 1e-18, 0.63, 0.646 \| PDLC00809, Cyclase_00049, 1.9e-18, 0.633, 0.722 |
| | B330r | CP124860 | cbass type II | QLQ95_17945 | PDLC00822 | Cyclase_00062 | Cyclase | 5.9999999999 | 6.8999999999 | 0.992 | 0.995 | 3248957 | 3250112 | + | - | 2841 | 4821 | PDLC00768.Cyclase_00008, 1, 0.0011, 0.411, 0.389 \| PDLC00769.Cyclase_00009, 1, 1.9e-11, 0.315, | PDLC00822, Cyclase_00062, 6.9e-161, 0.992, 0.995 \| PDLC00773, Cyclase_00013, 2.6e-42, 0.435, 0.5 \| PDLC00798, Cyclase_00038, 1e-19, 0.419, 0.471 \| PDLC00810, Cyclase_00050, 1.2e-18, 0.438, 0.525 \| PDLC00793, Cyclase_00033, 5.1e-17, 0.414, 0.348 |
| | B330r | CP124860 | cbass type II | QLQ95_17950 | PDLC00839 | E1E2_00015 | E1-E2 | 9.9999999999 | 1.2999999999 | 0.981 | 0.973 | 3250112 | 3251894 | + | - | 2842 | 4821 | PDLC00825.E1E2_00001, 1, 6.1e-11, 0.12, 0.125 \| PDLC00826.E1E2_00002, 1, 3.6e-17, 0.263, 0.27 | PDLC00839, E1E2_00015, 1.3e-232, 0.981, 0.973 \| PDLC00848, E1E2_00024, 6.3e-155, 0.978, 0.964 \| PDLC00835, E1E2_00011, 3.3e-86, 0.901, 0.695 \| PDLC00827, E1E2_00003, 1.3e-37, 0.605, 0.491 \| PDLC00852, E1E2_00028, 4.6e-37, 0.631, 0.647 |
| | B330r | CP124860 | cbass type II | QLQ95_17955 | PDLC00890 | JAB_00004 | JAB | 1.4e-52 | 1.5999999999 | 0.95 | 0.981 | 3251890 | 3252376 | + | - | 2843 | 4821 | PDLC00727.Effector_00065, 1, 4.5e-06, 0.609, 0.163 \| PDLC00827.E1E2_00003, 1, 8e-18, 0.894, 0.2 | PDLC00890, JAB_00004, 1.6e-52, 0.95, 0.981 \| PDLC00889, JAB_00003, 5.6e-50, 0.832, 0.81 \| PDLC00896, JAB_00010, 5.7e-25, 0.845, 0.849 \| PDLC00835, E1E2_00011, 2.2e-23, 0.826, 0.174 \| PDLC00902, JAB_00016, 3.3e-23, 0.913, 0.912 |
| | B330r | CP124860 | ietAS | QLQ95_14300 | PDLC02196 | IetA_00002 | IetA | 1.9999999999 | 2.1999999999 | 0.907 | 0.845 | 2466359 | 2467421 | - | - | 2121 | 4821 | PDLC02195.IetA_00001, 1, 1.7e-58, 0.918, 0.838 \| PDLC02196.IetA_00002, 1, 2.2e-72, 0.907, 0.845 \| | PDLC02196, IetA_00002, 2.2e-72, 0.907, 0.845 \| PDLC04232, IetA_WP_225151863.1, 8.1e-61, 0.918, 0.952 \| PDLC02195, IetA_00001, 1.7e-58, 0.918, 0.838 \| PDLC04231, IetA_WP_229193352.1, 4.8e-33, 0.731, 0.95 |
| | B330r | CP124860 | ietAS | QLQ95_14295 | PDLC02199 | IetS_00003 | IetS | 2,00E-36 | 3.4e-36 | 0.487 | 0.437 | 2463918 | 2466354 | + | - | 2120 | 4821 | PDLC02197.IetS_00001, 1, 5.9e-22, 0.307, 0.336 \| PDLC02197.IetS_00001, 2, 1.4e-12, 0.276, 0.271 \| | PDLC02199, IetS_00003, 3.4e-36, 0.487, 0.437 \| PDLC02198, IetS_00002, 8.8e-29, 0.531, 0.547 \| PDLC02201, IetS_00005, 1.1e-22, 0.483, 0.484 \| PDLC02197, IetS_00001, 5.9e-22, 0.307, 0.336 \| PDLC04233, IetS_WP_226782225.1, 1.4e-14, 0.402, 0.408 |
| | B330r | CP124860 | retron_VI | QLQ95_09720 | PDLC02926 | retron_type_VI_hth_cluster2_1 | HTH_VI | 8.5e-30 | 9.4e-30 | 0.929 | 0.978 | 1367061 | 1367358 | + | - | 1230 | 4821 | PDLC02926,retron_type_VI_hth_cluster2_1, 1, 9.4e-30, 0.929, 0.978 | PDLC02926, retron_type_VI_hth_cluster2_1, 9.4e-30, 0.929, 0.978 |
| | B330r | CP124860 | retron_VI | QLQ95_09725 | PDLC02952 | retron_VI | RT_VI | 1.1999999999 | 1.2999999999 | 0.929 | 0.885 | 1367332 | 1368175 | + | - | 1231 | 4821 | PDLC02952,retron_VI, 1, 1.3e-73, 0.929, 0.885 | PDLC02952, retron_VI, 1.3e-73, 0.929, 0.885 |
| | B330r | CP124860 | retron_VI | QLQ95_09715 | PDLC02970 | retron_type_VI_sp_cluster2760_1 | SP_VI | 2.0999999999 | 2.3e-30 | 0.932 | 0.971 | 1366771 | 1366996 | + | - | 1229 | 4821 | PDLC02970,retron_type_VI_sp_cluster2760_1, 1, 2.3e-30, 0.932, 0.971 | PDLC02970, retron_type_VI_sp_cluster2760_1, 2.3e-30, 0.932, 0.971 |
| | B330r | CP124860 | RM_type_I | QLQ95_14230 | PDLC03022 | MTase_I_00005 | MTase_I | 9.7999999999 | 1.1999999999 | 0.927 | 0.565 | 2447103 | 2448666 | - | - | 2108 | 4821 | PDLC03018.MTase_I_00001, 1, 9.7e-07, 0.165, 0.105 \| PDLC03018.MTase_I_00001, 2, 1.2e-05, 0.13 | PDLC03022, MTase_I_00005, 1.2e-167, 0.927, 0.565 \| PDLC03027, MTase_I_00010, 1.9e-110, 0.658, 0.43 \| PDLC03031, MTase_I_00014, 2.4e-97, 0.913, 0.582 \| PDLC03032, MTase_I_00015, 4.8e-85, 0.656, 0.349 \| PDLC03020, MTase_I_00003, 1.7e-74, 0.746, 0.951 |
| | B330r | CP124860 | RM_type_I | QLQ95_14210 | PDLC03048 | REase_I_00009 | REase_I | 2.8999999999 | 3.5999999999 | 0.949 | 0.944 | 2441702 | 2444582 | - | - | 2104 | 4821 | PDLC03040.REase_I_00001, 1, 3.8e-53, 0.757, 0.709 \| PDLC03041.REase_I_00002, 1, 0.049, 0.093, | PDLC03048, REase_I_00009, 3.6e-290, 0.949, 0.944 \| PDLC03046, REase_I_00007, 4.8e-73, 0.838, 0.846 \| PDLC03045, REase_I_00006, 1.3e-69, 0.6, 0.574 \| PDLC03040, REase_I_00001, 3.8e-53, 0.757, 0.709 \| PDLC03043, REase_I_00004, 1.3e-49, 0.268, 0.847 |
| | B330r | CP124860 | RM_type_I | QLQ95_14225 | PDLC03087 | Specificity_I_00036 | Specificity_I | 1.6999999999 | 1.9999999999 | 0.966 | 0.981 | 2445877 | 2447104 | - | - | 2107 | 4821 | PDLC03032.MTase_I_00015, 1, 6.5e-08, 0.348, 0.144 \| PDLC03032.MTase_I_00015, 2, 0.55, 0.228, 0 | PDLC03087, Specificity_I_00036, 2e-81, 0.966, 0.981 \| PDLC03178, Specificity_I_00127, 9.2e-70, 0.978, 0.993 \| PDLC03165, Specificity_I_00114, 1.5e-65, 0.975, 0.988 \| PDLC03085, Specificity_I_00034, 1.1e-47, 0.471, 0.468 \| PDLC03185, Specificity_I_00134, 4.6e-37, 0.975, 0.995 |
| | B330r | CP124860 | RM_type_II | QLQ95_10055 | PDLC03272 | MTase_II_00058 | MTase_II | 8.7999999999 | 9.9999999999 | 0.936 | 0.953 | 1440389 | 1441655 | - | - | 1296 | 4821 | PDLC03217.MTase_II_00003, 1, 4.1e-77, 0.767, 0.452 \| PDLC03220.MTase_II_00006, 1, 5.2e-31, 0.3 | PDLC03272, MTase_II_00058, 1e-130, 0.936, 0.953 \| PDLC03361, MTase_II_00147, 1.9e-101, 0.876, 0.866 \| PDLC03217, MTase_II_00003, 4.1e-77, 0.767, 0.452 \| PDLC03242, MTase_II_00028, 5.4e-71, 0.751, 0.716 \| PDLC03378, MTase_II_00164, 1.4e-65, 0.767, 0.956 |
| | B330r | CP124860 | VSPR | QLQ95_06595 | PDLC03272 | MTase_II_00058 | MTase_II | 2.5999999999 | 2.8999999999 | 0.979 | 0.801 | 651702 | 652689 | + | - | 615 | 4821 | PDLC03217.MTase_II_00003, 1, 7.1e-56, 0.927, 0.441 \| PDLC03220.MTase_II_00006, 1, 2.8e-20, 0.4 | PDLC03272, MTase_II_00058, 2.9e-111, 0.979, 0.801 \| PDLC03378, MTase_II_00164, 6.3e-69, 0.976, 0.98 \| PDLC03361, MTase_II_00147, 5.1e-62, 0.963, 0.785 \| PDLC03217, MTase_II_00003, 7.1e-56, 0.927, 0.441 \| PDLC03242, MTase_II_00028, 6.2e-44, 0.966, 0.746 |
| | B330r | CP124860 | VSPR | QLQ95_06600 | PDLC03397 | NEase_II_00006 | Vsr | 1.5e-12 | 1.8e-12 | 0.784 | 0.692 | 652685 | 653105 | - | - | 616 | 4821 | PDLC03397,NEase_II_00006, 1, 1.8e-12, 0.784, 0.692 | PDLC03397, NEase_II_00006, 1.8e-12, 0.784, 0.692 |
| | B330r | CP124860 | DMS_other | QLQ95_16095 | PDLC03398 | NEase_II_00007 | REase_II | 2.1e-16 | 2.7e-16 | 0.604 | 0.606 | 2868441 | 2869110 | - | - | 2477 | 4821 | PDLC03398,NEase_II_00007, 1, 2.7e-16, 0.604, 0.606 | PDLC03398, NEase_II_00007, 2.7e-16, 0.604, 0.606 |
| | B330r | CP124860 | RM_type_II | QLQ95_10050 | PDLC03563 | REase_II_00165 | REase_II | 5.4999999999 | 6.0999999999 | 0.994 | 0.984 | 1439313 | 1440390 | - | - | 1295 | 4821 | PDLC03563,REase_II_00165, 1, 6.1e-113, 0.994, 0.984 | PDLC03563, REase_II_00165, 6.1e-113, 0.994, 0.984 |
| | B330r | CP124860 | RM_type_IV | QLQ95_12110 | PDLC03657 | mREase_IV_00003 | mREase_IV | 1.9e-60 | 2.9e-60 | 0.977 | 0.991 | 1928385 | 1929309 | - | - | 1701 | 4821 | PDLC03657.mREase_IV_00003, 1, 2.9e-60, 0.977, 0.991 \| PDLC03663.mREase_IV_00009, 1, 7e-11, | PDLC03657, mREase_IV_00003, 2.9e-60, 0.977, 0.991 \| PDLC03663, mREase_IV_00009, 7e-11, 0.407, 0.211 |
| | B330r | CP124860 | SoFic | QLQ95_14705 | PDLC03963 | SoFic__SoFic | SoFic | 8.8e-49 | 1.1999999999 | 0.728 | 0.839 | 2557022 | 2558315 | - | - | 2199 | 4821 | PDLC03963,SoFic__SoFic, 1, 1.2e-48, 0.728, 0.839 | PDLC03963, SoFic__SoFic, 1.2e-48, 0.728, 0.839 |
| | B330r | CP124860 | DMS_other | QLQ95_16075 | PDLC04135 | mads_3 | mad3 | 1.4999999999 | 1.2e-41 | 0.345 | 0.971 | 2862673 | 2865469 | - | - | 2473 | 4821 | PDLC04135.mads_3, 1, 8.8e-13, 0.098, 0.267 \| PDLC04135.mads_3, 2, 4.7e-27, 0.223, 0.572 \| PDLC | PDLC04135, mads_3, 1.2e-41, 0.345, 0.971 \| PDLC04140, mads_8, 1.1e-12, 0.302, 0.732 |
| | B330r | CP124860 | PDC-M15 | QLQ95_14220 | PDLC04455 | PDC-M15A_WP_018278665.1 | PDC-M15A | 2.6999999999 | 3.1999999999 | 0.969 | 0.973 | 2445182 | 2445863 | - | - | 2106 | 4821 | PDLC04454.PDC-M15A_WP_123962630.1, 1, 4.8e-40, 0.735, 0.786 \| PDLC04455.PDC-M15A_WP_0 | PDLC04455, PDC-M15A_WP_018278665.1, 3.2e-84, 0.969, 0.973 \| PDLC04454, PDC-M15A_WP_123962630.1, 4.8e-40, 0.735, 0.786 \| PDLC04456, PDC-M15A_WP_017232921.1, 2e-32, 0.836, 0.696 \| PDLC04458, PDC-M15A_WP_220777606.1, 8.5e-31, 0.885, 0.848 \| PDLC04459, PDC-M15A_WP_199528910.1, 4.5e-28, 0.85, 0.797 |
| | B330r | CP124860 | PDC-M15 | QLQ95_14215 | PDLC04463 | PDC-M15B_WP_231555654.1 | PDC-M15B | 6.5999999999 | 7.2e-67 | 0.955 | 0.942 | 2444596 | 2445202 | - | - | 2105 | 4821 | PDLC04460.PDC-M15B_WP_061013001.1, 1, 2.8e-08, 0.886, 0.91 \| PDLC04461.PDC-M15B_WP_12 | PDLC04463, PDC-M15B_WP_231555654.1, 7.2e-67, 0.955, 0.942 \| PDLC04468, PDC-M15B_WP_132075670.1, 8.1e-16, 0.632, 0.626 \| PDLC04460, PDC-M15B_WP_061013001.1, 2.8e-08, 0.886, 0.91 \| PDLC04462, PDC-M15B_WP_011787717.1, 5.5e-07, 0.632, 0.626 \| PDLC04465, PDC-M15B_WP_220777607.1, 1.3e-06, 0.915, 0.908 |
| | B330r | CP124860 | PDC-S02 | QLQ95_14325 | PDLC04822 | PDC-S02_WP_207168644.1 | PDC-S02 | 1.3999999999 | 1.6999999999 | 0.841 | 0.976 | 2475001 | 2476024 | + | - | 2126 | 4821 | PDLC04793.PDC-S02_WP_149946094.1, 1, 9.6e-20, 0.224, 0.658 \| PDLC04794.PDC-S02_WP_0912 | PDLC04822, PDC-S02_WP_207168644.1, 1.7e-135, 0.841, 0.976 \| PDLC04807, PDC-S02_WP_151674975.1, 9.1e-114, 0.95, 0.954 \| PDLC04795, PDC-S02_WP_011937941.1, 1.2e-82, 0.926, 0.899 \| PDLC04831, PDC-S02_WP_143549780.1, 3.5e-80, 0.909, 0.939 \| PDLC04810, PDC-S02_WP_080642564.1, 3e-77, 0.932, 0.935 |
| | B330r | CP124860 | PDC-S06 | QLQ95_01430 | PDLC04890 | PDC-S06_WP_095244428.1 | PDC-S06 | 2.4999999999 | 2.7999999999 | 0.999 | 0.986 | 5228709 | 5230974 | - | - | 4434 | 4821 | PDLC04886.PDC-S06_WP_029531116.1, 1, 1.8e-129, 0.999, 0.993 \| PDLC04887.PDC-S06_WP_221 | PDLC04890, PDC-S06_WP_095244428.1, 2.8e-168, 0.999, 0.986 \| PDLC04886, PDC-S06_WP_029531116.1, 1.8e-129, 0.999, 0.993 \| PDLC04897, PDC-S06_WP_115337947.1, 4e-57, 0.995, 0.99 \| PDLC04894, PDC-S06_WP_100093761.1, 3.7e-39, 0.984, 0.972 \| PDLC04887, PDC-S06_WP_221405099.1, 8.8e-38, 0.874, 0.858 |
| | B330r | CP124860 | PDC-S13 | QLQ95_17535 | PDLC04919 | PDC-S13_WP_186842295.1 | PDC-S13 | 2.6999999999 | 3.0999999999 | 0.965 | 0.968 | 3170396 | 3171266 | + | - | 2762 | 4821 | PDLC04918.PDC-S13_WP_217993064.1, 1, 1.4e-07, 0.204, 0.276 \| PDLC04918.PDC-S13_WP_2179 | PDLC04919, PDC-S13_WP_186842295.1, 3.1e-92, 0.965, 0.968 \| PDLC04918, PDC-S13_WP_217993064.1, 2.4e-34, 0.408, 0.556 \| PDLC04920, PDC-S13_WP_097300085.1, 1.1e-12, 0.384, 0.392 |
| | B330r | CP124860 | PDC-S38 | QLQ95_23745 | PDLC05010 | PDC-S38_WP_174243465.1 | PDC-S38 | 1.9999999999 | 2.1e-31 | 0.982 | 0.982 | 4651236 | 4651410 | + | - | 3983 | 4821 | PDLC05010,PDC-S38_WP_174243465.1, 1, 2.1e-31, 0.982, 0.982 | PDLC05010, PDC-S38_WP_174243465.1, 2.1e-31, 0.982, 0.982 |
